# NOP16 is a histone mimetic that regulates Histone H3K27 methylation and gene repression

**DOI:** 10.1101/2023.06.13.544862

**Authors:** Ken Takashima, Dian-Jang Lee, María Fernanda Trovero, M. Hafiz Rothi, Meeta Mistry, Ying Zhang, Zhouyihan Li, Christopher P. Davis, Zilan Li, Julia Natale, Ernst Schmid, Joseph Al Haddad, Gabriela Brunsting Hoffmann, Sabine Dietmann, Shannan Ho Sui, Hiroyuki Oshiumi, Judy Lieberman, Eric Lieberman Greer

**Affiliations:** Department of Pediatrics, HMS Initiative for RNA Medicine, Harvard Medical School, Boston MA, USA; Division of Newborn Medicine, Boston Children’s Hospital, Boston MA, USA; Department of Immunology, Graduate School of Medical Sciences, Faculty of Life Sciences, Kumamoto University, Kumamoto, Japan; Program in Cellular and Molecular Medicine, Boston Children’s Hospital, Boston MA, USA; Bioinformatics Core, Department of Biostatistics, Harvard T.H. Chan School of Public Health, Boston MA, USA; Department of Neurobiology, Harvard Medical School, Boston MA, USA; Department of Developmental Biology and Center of Regenerative Medicine, Washington University School of Medicine, St. Louis, MO 63110, USA; Department of Pediatrics, Washington University School of Medicine, St. Louis, MO 63110, USA; Department of Genetics, Washington University School of Medicine, St. Louis, MO 63110, USA

**Keywords:** H3K27me3, histone mimic, NOP16, breast cancer, PRC2, JMJD3, EED

## Abstract

Post-translational modifications of histone tails alter chromatin accessibility to regulate gene expression. Some viruses exploit the importance of histone modifications by expressing histone mimetic proteins that contain histone-like sequences to sequester complexes that recognize modified histones. Here we identify an evolutionarily conserved and ubiquitously expressed, endogenous mammalian protein Nucleolar protein 16 (NOP16) that functions as a H3K27 mimic. NOP16 binds to EED in the H3K27 trimethylation PRC2 complex and to the H3K27 demethylase JMJD3. NOP16 knockout selectively globally increases H3K27me3, a heterochromatin mark, without altering methylation of H3K4, H3K9, or H3K36 or acetylation of H3K27. NOP16 is overexpressed and linked to poor prognosis in breast cancer. Depletion of NOP16 in breast cancer cell lines causes cell cycle arrest, decreases cell proliferation and selectively decreases expression of E2F target genes and of genes involved in cell cycle, growth and apoptosis. Conversely, ectopic NOP16 expression in triple negative breast cancer cell lines increases cell proliferation, cell migration and invasivity *in vitro* and tumor growth *in vivo*, while NOP16 knockout or knockdown has the opposite effect. Thus, NOP16 is a histone mimic that competes with Histone H3 for H3K27 methylation and demethylation. When it is overexpressed in cancer, it derepresses genes that promote cell cycle progression to augment breast cancer growth.

## Introduction

Trimethylation of H3K27 is a hallmark of inactive gene promoters and heterochromatin. H3K27 is methylated by the polycomb repressive complex 2 (PRC2), comprised of EZH2, EED, SUZ12 and other components ^1^, and is demethylated by UTX or JMJD3 ^2, 3^. In some cancers, including breast cancer, EZH2 is overexpressed, and overexpression has been linked to increased cell proliferation and tumor malignancy ^4^. While EZH2 overexpression causes increased H3K27me3 in B-cell lymphomas, EZH2 overexpression in breast cancers does not correlate with elevated H3K27me3 ^5^, suggesting that additional mechanisms that regulate H3K27me3 remain to be discovered. Histone methyltransferases can also methylate non-histone molecules ^6, 7^. Several non-histone viral targets of histone methyltransferases, such as G9a and influenza A NS1 protein, contain short amino acid sequences homologous to histone tails ^8, 9^. Some viruses contain ∼4 amino acid histone-like sequences that can interfere with epigenetic regulation in host cells to regulate the host immune response to viral infection ^10–12^. Endogenous histone mimetic proteins that might similarly regulate host gene expression have not been described.

In this study, we searched for mammalian proteins containing sequences homologous to the region surrounding H3K27. Nucleolar protein 16 (NOP16) stood out because of its high homology in the region surrounding K29 (ARRK29AAP in NOP16 vs AARK27SAP in histone H3) (Fig. 1a). NOP16 is a poorly characterized, ubiquitously expressed, 178 amino acid nuclear protein encoded on human chromosome 5q35.2 with orthologs throughout the Chordata (Extended Data Fig. 1a). NOP16 has been implicated in regulating rRNA biogenesis ^13^. It is overexpressed in some cancers and its expression has been linked to higher cell proliferation and poor prognosis in breast cancer ^13–15^. In breast cancer both estrogen and c-Myc strongly enhance *Nop16* transcription ^14^. Of note, *Nop16* has turned up as a “hit” that affects cell proliferation in CRISPR gene knockout screens in multiple cancer cell lines ^16^. However, whether or how it increases cell proliferation is uncertain. We found that NOP16 is expressed similarly in a number of human cancer cell lines including representative breast cancer cell lines MCF7, MDA-MB231, and MDA-MB468 (Extended Data Fig. 1b). Here we show that NOP16 is methylated, binds to both EED in PRC2 and JMJD3, and NOP16 knockout selectively increases H3K27me3, suggesting that NOP16 is a histone H3 mimetic that competes with histone H3 for binding to the H3 methylation/demethylation machinery. In breast cancer cell lines, deficiency of NOP16 suppresses tumor cell proliferation, cell cycle progression, migration and invasivity *in vitro* and tumor cell growth *in vivo* by selectively increasing H3K27me3 especially of target genes of the E2F transcription factors, key regulators of the G1-> S transition, and other genes that control cell proliferation and cell death.

**Figure 1.**
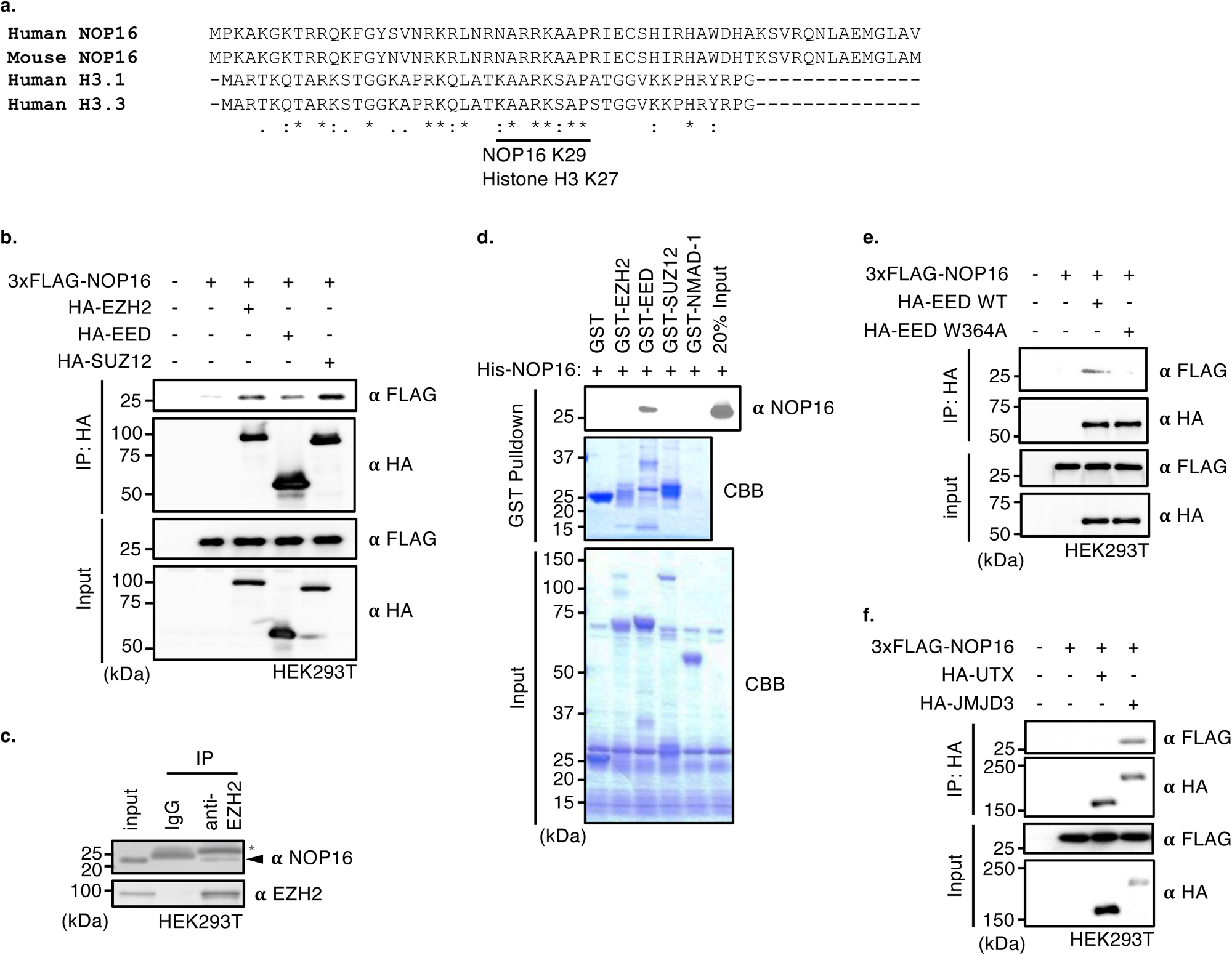
NOP16 interacts with H3K27me3 modifiers. **a.** The N terminus of NOP16 displays high similarity to the histone H3 tail. Alignment of NOP16 N-terminal region and histone H3 tails. “*”, “:”, and “.” indicate identical, strong similarity, and weak similarity, respectively. **b.** NOP16 binds to PRC2 components *ex vivo*. 3xFLAG-tagged NOP16 and/or HA-tagged PRC2 complex (EZH2, EED, or SUZ12) expressing vectors were transfected into HEK293T cells. After immunoprecipitation with an HA antibody, interaction with NOP16 was assessed by western blotting. **c.** Endogenous NOP16 binds to EZH2. Immunoprecipitations with antibodies directed against EZH2 or control IgG were performed and binding was assessed by western blotting using anti-NOP16 antibody. **d.** NOP16 directly binds to EED as assessed by *in vitro* binding assays. 6xHis-tagged NOP16 was incubated with GST, GST-tagged EZH2, GST-tagged EED, GST-tagged SUZ12 or GST-tagged NMAD-1. After GST pull-down, binding was assessed by western blotting with a His antibody. **e.** EED WD40 domain is required for binding to NOP16 HEK293T cells were transfected with 3xFLAG-tagged NOP16 and/or HA-tagged EED wildtype (WT) or WD40 domain mutant W364A, proteins were immunoprecipitated with an HA antibody, and interactions were detected by western blotting. **f.** NOP16 binds to histone demethylase JMJD3 not but UTX *ex vivo*. HEK293T cells transfected with 3xFLAG-tagged NOP16 and/or HA-tagged UTX or JMJD3, immunoprecipitated with an HA antibody, and subjected to western blotting with HA and Flag antibodies to detect binding.

## Results

### NOP16 interacts with H3K27 modifiers

We performed a BLAST search to identify host proteins containing sequence homology to the histone H3 tail and identified NOP16 (Fig. 1a). Because of the high sequence similarity of NOP16 to the histone H3 tail around lysine 29 of NOP16 and lysine 27 of H3, we hypothesized that NOP16 could function as a histone tail mimic. To assess this possibility, we examined whether NOP16 physically interacted with H3K27 modifiers. Ectopically expressed 3xFLAG-tagged NOP16 co-immunoprecipitated with HA-tagged components of the polycomb repressive complex 2 (PRC2), including the methyltransferase EZH2, the zinc finger protein SUZ12, and the H3K27me3 binding protein EED expressed in HEK293T cells (Fig. 1b). Furthermore, endogenous NOP16 co-immunoprecipitated with endogenous EZH2 (Fig. 1c), suggesting that this interaction was not an overexpression artefact. Cell fractionation revealed that both NOP16 and EZH2 are present in both chromatin-bound and chromatin-free nuclear fractions (Extended Data Fig. 1c). To identify which component of the PRC2 complex directly interacted with NOP16, we performed an *in vitro* binding assay using His-tagged NOP16 and glutathione S-transferase (GST)-tagged EZH2, EED, and SUZ12 recombinant proteins that were expressed and purified from *E. coli*. NOP16 selectively bound to GST-tagged EED but not to GST, other components of the PRC2 complex, or a control protein ^17^ (Fig. 1d). Together, these data indicate that NOP16 directly interacts with EED.

EED binds to H3K27me3 modified histones via its WD40 domain ^18^, which recruits the PRC2 complex to target genomic regions and sustains propagative H3K27me3 ^18^. To test whether the WD40 domain of EED is important for binding to NOP16, we expressed WT and a W364A mutant of EED, which disrupts WD40 domain binding ^18^, in HEK293T cells. Disruption of the WD40 domain of HA-tagged EED prevented binding to NOP16 *ex vivo* (Fig. 1e), indicating that the WD40 domain of EED mediates binding to both NOP16 and histone H3. GST-tagged NOP16 truncation mutants were used to identify the region of NOP16 responsible for EED binding. Both NOP16 1-38 and 39-176 pulled down EED (Extended Data Fig. 1d), suggesting that NOP16 contains multiple EED binding sites. Shorter truncations of NOP16 (amino acids 1-22, 23-44, and 67-88) also pulled down EED with varying efficiency, but the N-terminal 1-22 peptide was most efficient and bound comparably to full-length NOP16 (Extended Data Fig. 1e). These data suggest that NOP16 has multiple interactions with EED.

To determine whether NOP16 also interacts with an H3K27me3 demethylase we performed immunoprecipitation assays with HA-tagged JMJD3 and UTX, the two known H3K27me3 demethylases. NOP16 interacted with HA-tagged JMJD3, but not with UTX (Fig. 1f). An *in vitro* binding assay revealed that His-tagged NOP16 directly bound to the JMJD3 JmjC domain (Extended Data Fig. 1f). Thus NOP16 interacts directly with both the H3K27me3 demethylase JMJD3 and EED in the H3K27 tri-methyltransferase complex.

### Methylation of NOP16 affects the binding affinity between NOP16 and EED

To determine whether NOP16 is methylated, we immunoprecipitated NOP16 from HEK293T cells expressing FLAG-tagged NOP16 and probed western blots with a pan-methyl lysine antibody. The pan-methyl lysine antibody detected a specific band (Fig. 2a), suggesting that NOP16 is methylated in cells. To determine which residue of NOP16 was methylated, we immunoprecipitated NOP16 and performed mass spectrometry, which suggested that either lysine 106 or 107 were methylated (Extended Data Fig. 2a). While lysine 29, which showed the closest homology to H3K27 was not detected by mass spectrometry this does not exclude that this residue might be methylated as no peptides containing lysine 29 were detected in independent mass spectrometry experiments. Substitution of both lysines 106 and 107 with alanines (K106A K107A) reduced NOP16 methylation as detected by pan-methyl lysine antibody in cells, but the K29A mutation did not (Fig. 2b). To assess the physiological significance of NOP16 methylation, we investigated whether the methylated lysines were important for NOP16 binding to EED. While substitutions of K29 or K106 and K107 of NOP16 with alanines did not reduce NOP16 binding to EED, substitution of all three lysines (K29, K106, and K107) strongly attenuated EED binding (Fig. 2c). These results suggest that methylation of all these lysines regulates NOP16’s interaction with EED. *In vitro* pulldown of a short synthetic NOP16 aa 25-32 peptide containing unmethylated or mono-, di- or tri-methylated K29 with EED showed that only the peptides containing di- or tri-methylated K29 bound to GST-EED (Extended Data Fig. 2b). To assess the affinity of EED for differently methylated NOP16 fragments we performed microscale thermophoresis ^19, 20^ with methylated NOP16 and histone tail peptides. EED bound most tightly and similarly to H3K27me3 and NOP16_25-32_K29me3 peptides (K_D_ = 0.95 ± 0.84 μM and K_D_ = 1.21 ± 1.07 μM, respectively, p = 0.9891 by one-way ANOVA), but had significantly lower affinity for un-, mono-, or dimethylated K29 (K_D_ = 11.64 ± 17.67 μM, 100.25 ± 77.16 μM, and 131.33 ± 14.57 μM, respectively) (Figure 2D). On the other hand, *in vitro* binding pulldown of JMJD3 JmjC domain was unaffected by NOP16 K29 methylation (Extended Data Fig. 2c). Similar to EED, JMJD3 JmjC domain bound most strongly to the H3K27me3 peptide (K_D_ = 8.59 ± 4.54 μM) and NOP16_25-32_K29me3 peptides (K_D_ = 32.15 ± 27.37 μM, p = 0.4303 by one-way ANOVA), and had significantly lower affinity for un-, mono-, or dimethylated K29 (K_D_ = 102.68 ± 65.25 μM, 346.67 ± 224.26 μM, and 305.6 ± 100.8 μM, respectively) as assessed by microscale thermophoresis (Extended Data Fig. 2d). Furthermore, the GST-tagged EED W364A WD40 domain mutant failed to bind to NOP16 trimethylated K29 peptides (Fig. 2e), indicating that methylated K29 NOP16 interacts with EED through EED’s WD40 domain. Thus, NOP16 is methylated and methylation is required for NOP16 binding to EED.

**Figure 2.**
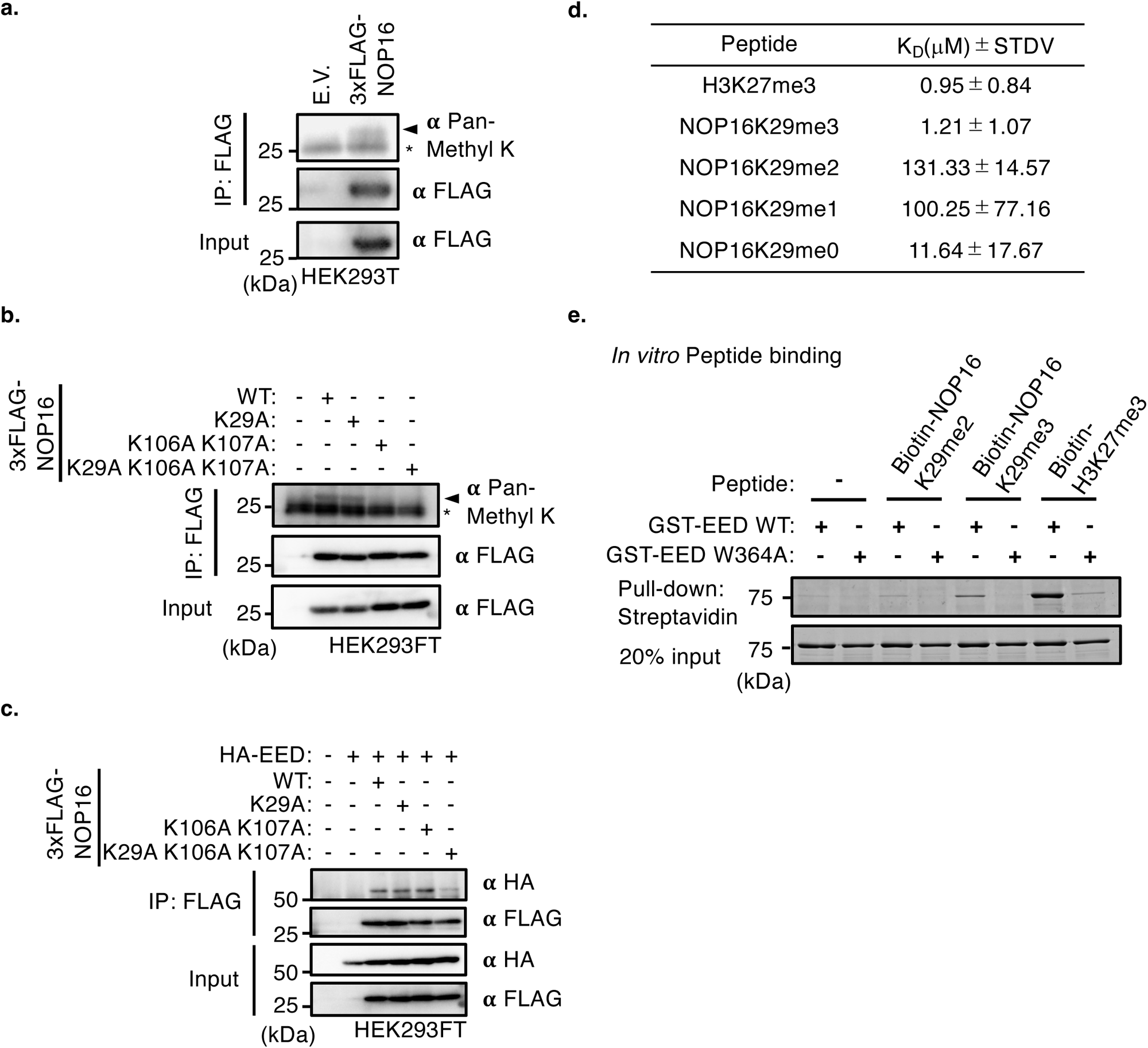
NOP16 methylation affects EED binding. **a.** NOP16 is methylated *ex vivo.* 3xFLAG-tagged NOP16 was immunoprecipitated from HEK293T cells and western blots were probed with a pan-methyl lysine antibody. **b.** Lysine 106 or 107 of NOP16 is methylated *ex vivo*. 3xFlag-tagged NOP16 wild-type (WT), lysine 29 to alanine (K29A), K106A and K107A double mutant, or K29A, K106A, and K107A triple mutant were transfected into HEK293FT cells, NOP16 was immunoprecipitated with a FLAG antibody, and methylation was assessed by western blotting using a pan-methyl lysine antibody. **c.** NOP16 lysines 29, 106, and 107 are important for binding to EED. Triple mutant 3xFlag-tagged NOP16 (K29A, K106A, and K107A) displayed reduced binding to HA-tagged EED as assessed by western blotting of immunoprecipitation reaction. **d.** Microscale thermophoresis of EED and methylated or unmethylated NOP16 or H3K27me3 peptides show that EED has highest affinity for H3K27me3, followed by NOP16K29me3, NOP16K29me0, NOP16 K29me1, and NOP16K29me2 in decreasing order of affinity. This table represents the mean +/- SD of 3-7 independent experiments. **e.** The WD40 domain of EED is necessary for binding to methylated NOP16 peptides. 1 μg of biotinylated synthetic peptides which were di(me2) or tri(me3)-methylated NOP16 amino acids 25-32 or H3K27me3 were incubated with 10 μg of GST-tagged WT or WD40 domain mutant (W364A) EED. Binding was detected by imperial staining after streptavidin pull-down.

### NOP16 negatively regulates H3K27me3

Since NOP16 binds to both an H3K27 methyltransferase complex and demethylase, we next assessed whether NOP16 affects histone H3 methylation. *NOP16* knockdown (KD) increased overall H3K27me3 in MCF7 and MDA-MB231 breast cancer cell lines without affecting levels of EZH2 (Fig. 3a and Extended Data Fig. 3a). Similarly, *NOP16* knockout by CRISPR-Cas9 also increased H3K27me3 in MDA-MB231 (Fig. 3b). However, knockout of NOP16 did not affect other histone modifications including H3K4me3, H3K9me3, H3K27Ac, or H3K36me3. Conversely, ectopic expression of *NOP16* in MDA-MB231 decreased overall H3K27me3 compared to empty vector (EV) control (Fig. 3c). These data indicate that NOP16 negatively regulates H3K27me3. To investigate whether NOP16 regulates H3K27me3 in specific genomic regions, we performed cleavage under targets and release using nuclease (CUT&RUN) ^21^ with an H3K27me3 specific antibody using MDA-MB231 cells. The levels and distribution of H3K27me3 were changed by *NOP16* over-expression at many sites scattered throughout the genome (Fig. 3d). In *NOP16* over-expressing cells, H3K27me3 was significantly changed at 1063 sites compared to control cells (Extended Data Fig. 3b). Of these, 755 sites showed decreased H3K27me3, and 308 sites had increased levels of H3K27me3 (FDR<0.05). Together, these results suggest that NOP16 inhibits H3K27 trimethylation.

**Figure 3.**
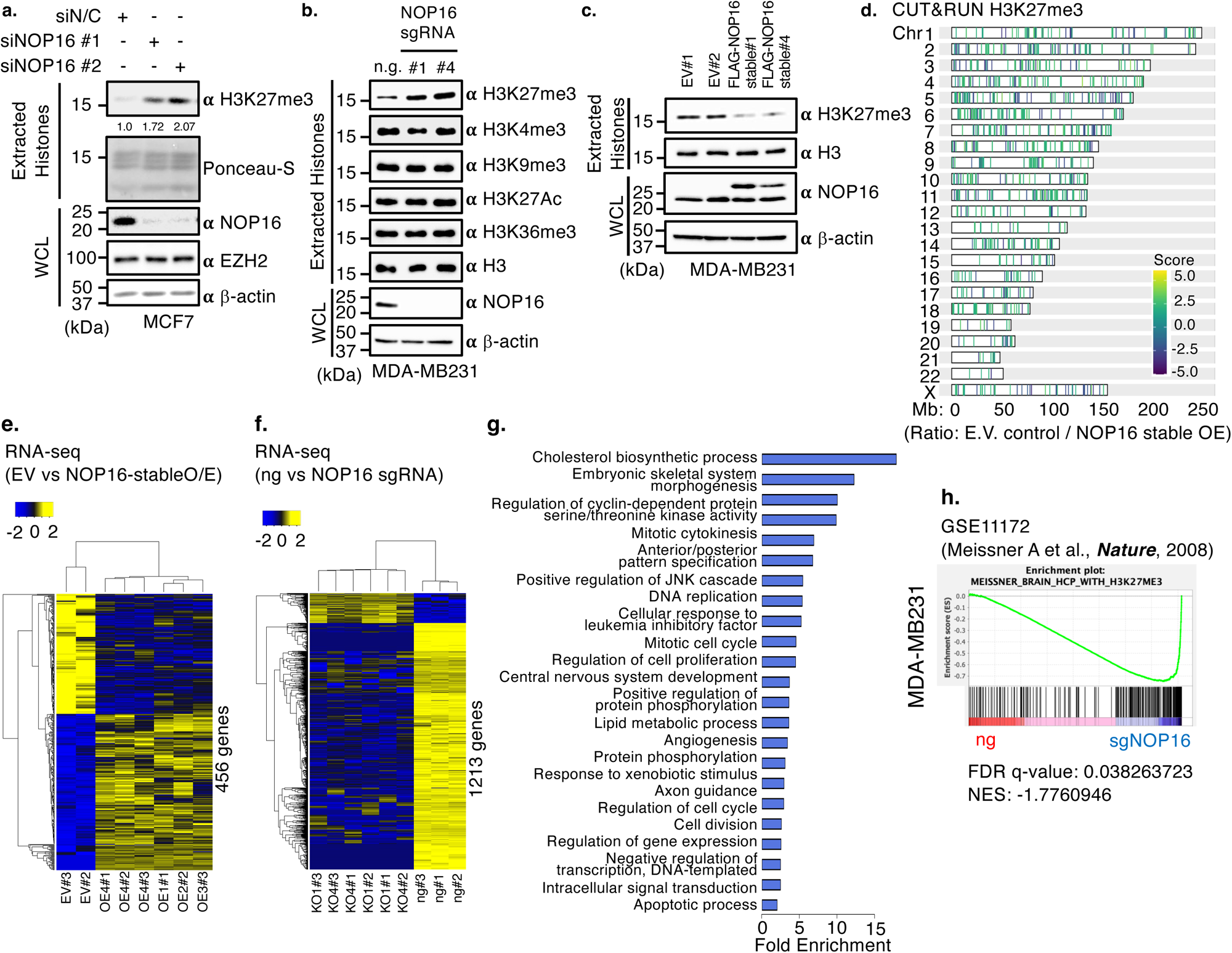
NOP16 negatively regulates H3K27me3. **a.** Knock down of NOP16 increases H3K27me3 in MCF7 breast cancer cells. MCF7 cells were transfected with negative control (N/C) or NOP16 siRNAs (10 pmol) and levels of H3K27me3 was determined by western blotting of extracted histones. Ponceau-S staining is used to examine total histone levels while NOP16, EZH2 and β-actin levels are shown by western blotting of whole cell lysates (WCL). **b.** NOP16 knockout causes a specific increase in H3K27me3. MDA-MB231 cells were transfected with Cas9 and infected with no guide (n.g.) or single guide RNAs against NOP16 (sgRNA #1 or #4) and extracted histones were assessed by western blotting for modifications. The efficiency of NOP16 deletion was assessed by western blotting of whole cell lysates. **c.** NOP16 stable-over expression reduces the global levels of H3K27me3 in MDA-MB231 cells relative to an empty vector control as assessed by western blotting of extracted histones. **d.** NOP16 stable overexpression decreased the levels of H3K27me3 across all chromosomes as assessed by CUT&RUN. Score and band color reflects the ratio of H3K27me3 intensity between E.V. control and NOP16 stably overexpressed MDA-MB231 cells. **e.** Heatmap of differentially expressed genes between NOP16 overexpression cells vs control cells. 258 up-regulated genes (padj<0.05, log2FC>=0.5) and 198 down-regulated genes (padj<0.05, log2FC<-0.5). **f.** Heatmap of differentially expressed genes between NOP16 knockout vs control MDA-MB231 cells. 131 up-regulated genes (padj<0.05, log2FC>1.5) and 1082 down-regulated genes (padj<0.05, log2FC<-1.5). **g.** Gene ontology analysis of downregulated genes in MDA-MB231 cells in response to NOP16 knockout and in Raw264.7 cells in response to Nop16 knockdown. **h.** NOP16 knockout induces changes in genes with high-CpG-density promoters (HCP) and H3K27me3 modified histones. This plot shows gene set enrichment analysis of transcripts that are differentially expressed after NOP16 knock out using GSE11172 gene sets. FDR: False Discovery Rate, NES: normalized enrichment score.

To determine whether NOP16 regulates gene expression, we performed RNA sequencing after overexpression or knockout of *NOP16* in MDA-MB231 cells. *NOP16* overexpression significantly up-regulated 258 genes and down-regulated 198 genes (Fig. 3e). Gene ontology (GO) analysis revealed that NOP16 overexpression reduced expression of genes involved in cell development and interaction with the extracellular environment (Extended Data Fig. 4a). *NOP16* knockout significantly up-regulated 131 genes and down-regulated 1082 genes (Fig. 3f). We used a more stringent cutoff to identify differentially expressed genes for the knockout than we used for ectopic *NOP16* overexpression since more genes were modulated by knockout, presumably because the cell line already expressed endogenous NOP16. To determine whether the consequences of manipulating NOP16 were conserved, we performed RNA sequencing after knockdown of Nop16 in mouse macrophage Raw264.7 cells. NOP16 knockdown significantly up-regulated 191 genes and down-regulated 3015 genes, confirming that NOP16 depletion, which causes an increase in the repressive chromatin modification H3K27me3, leads to decreased gene expression as expected. We found that 6 genes were consistently up-regulated and 314 genes were consistently down-regulated in both human MDA-MB231 cells and Raw264.7 cells in response to NOP16 depletion. A gene ontology analysis revealed that the most downregulated processes after NOP16 knockout or knockdown in human and mouse cells were involved in lipid metabolism, development, mitosis, DNA replication, cell cycle, cell proliferation, cell division, and apoptosis (Fig. 3g and Extended Data Fig. 4b-d). Interestingly and consistently, a previous study showed that NOP16 altered lipid metabolism purportedly by binding to ATP-citrate lyase, which catalyzes the production of acetyl-coA, which is needed for fatty acid and cholesterol biosynthesis, and indirectly increased H3K27 acetylation^22^. There was a small but statistically significant overlap of genes that were downregulated in response to NOP16 KO and genes that were upregulated in response to NOP16 overexpression (Extended Data Fig. 4e, 20/238; p<0.00001), while there was no-overlap of genes that were upregulated in NOP16 KO and genes that were downregulated in NOP16 overexpression in MDA-MB231 cells (Extended Data Fig. 4f). A gene set enrichment analysis (GSEA) revealed that genes downregulated in response to NOP16 deletion were enriched in genes with high-CpG-density promoters (HCP) that bear the H3K27me3 modification in various cell types or tissues (Fig. 3h and Extended Data Fig. 4g), suggesting that NOP16 regulates genes that are marked with H3K27me3. Disease-related GO analysis revealed that NOP16 overexpression or knockout in MDA-MB231 cells altered the expression of tumor-related genes in kidney, renal cell carcinoma, bile duct carcinoma (Extended Data Fig. 5a-c), and CNS tumors (Extended Data Fig. 5d-f). Moreover, integration analysis of CUT&RUN and RNA-seq data showed a small, but highly significant, overlap of genes that were up-regulated and had decreased H3K27me3 in response to *NOP16* overexpression (Extended Data Fig. 6, 21/237, p < 0.0001 by hypergeometric probability). Together these data suggest that NOP16 negatively regulates H3K27me3 and that overexpression of NOP16 and the subsequent decrease in H3K27me3 causes an increase in transcription of genes involved in cell cycle and tumorigenesis.

**Figure 4.**
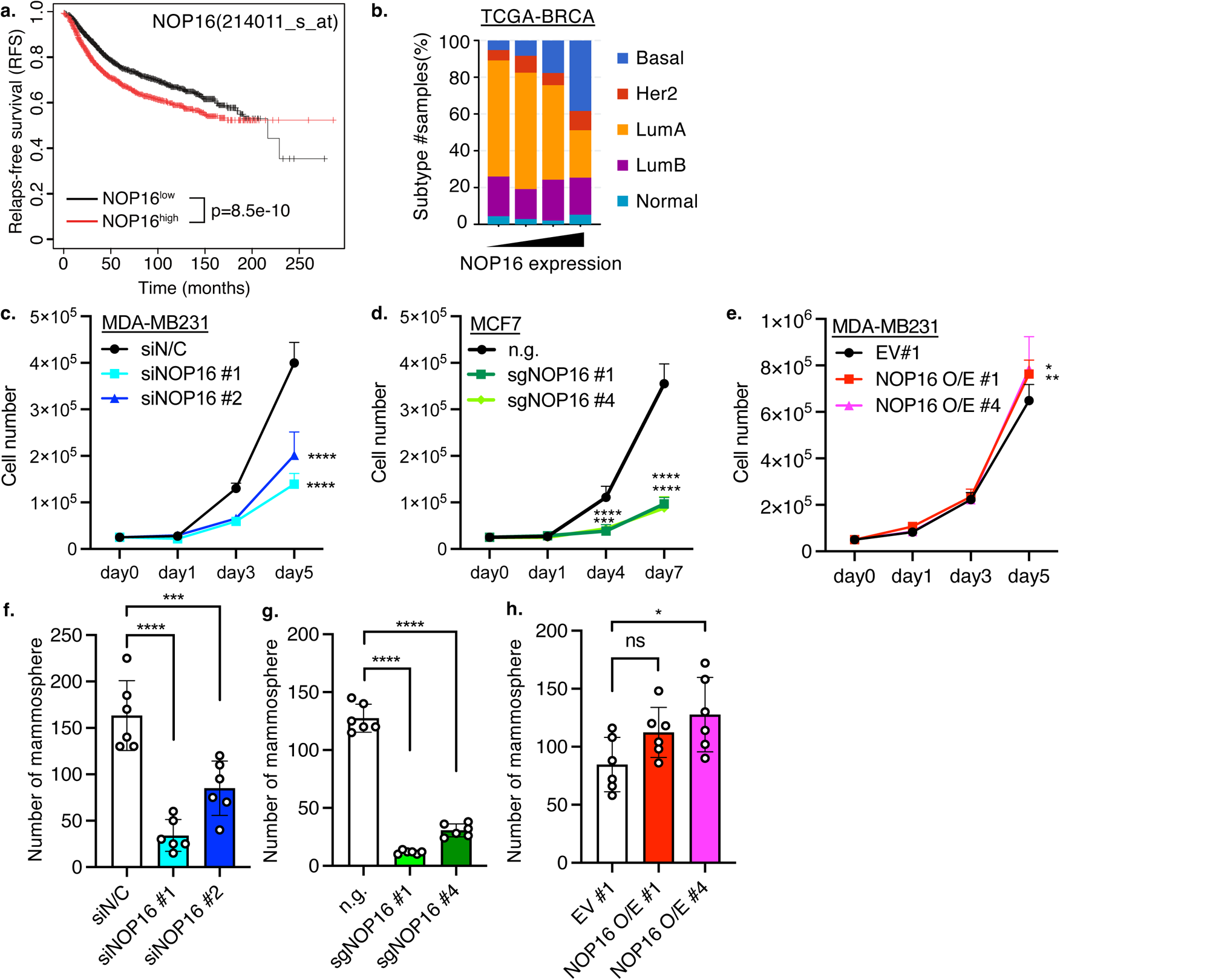
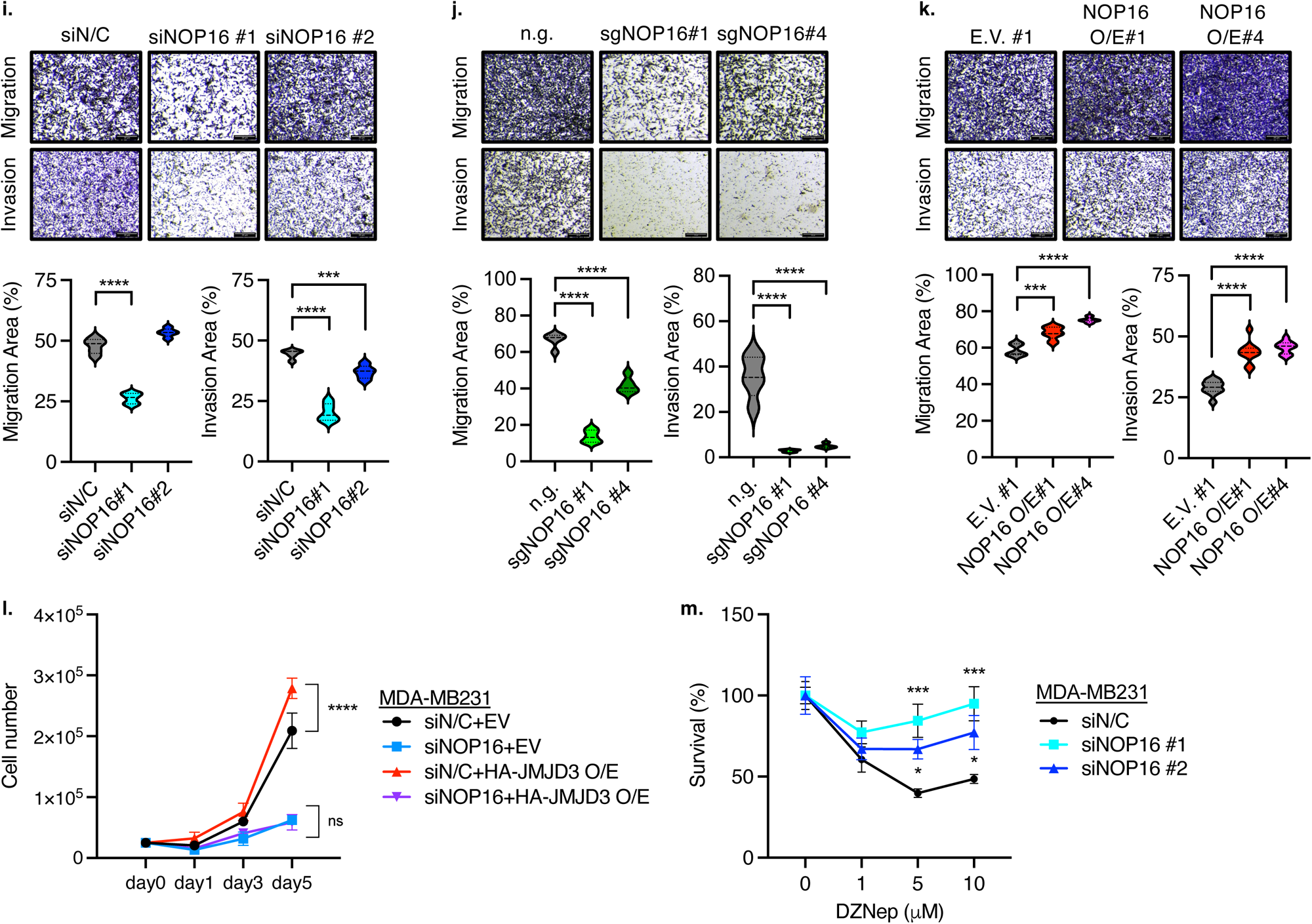
NOP16 maintains cell proliferation and migration of breast cancer cells. **a.** Higher expression of NOP16 correlates with poorer prognosis in breast cancer patients as visualized from Kaplan-Meier plotter. **b.** Elevated expression of NOP16 correlates with a relative increase in basal breast cancer subtype. TCGA-BRCA data sets were acquired from cBioPortal for Cancer Genomics. **c.** NOP16 knockdown in MDA-MB231 cells decreases proliferation rates relative to a negative control (N/C) knockdown. ****p<0.001 as assessed by two-way ANOVA followed by Dunnett’s post-test. **d.** NOP16 knockout slows cell proliferation in MCF7 cells. MCF7 cells were infected with lentivirus encoding Cas9 and single guide RNAs directed against control (n.g.) or NOP16 (#1 or #4). ****p<0.001 as assessed by two-way ANOVA followed by Dunnett’s post-test. **e.** NOP16 overexpression causes a subtle increase in cell growth of MDA-MB231 breast cancer cells. Proliferation of MDA-MB231 stably overexpressing 3xFLAG-NOP16 (#1 or #4) was assessed relative to an empty vector (EV) control line. *p<0.05, **p<0.01 as assessed by two-way ANOVA followed by Dunnett’s post-test. **f-h.** NOP16 is essential for scaffold-independent mammosphere formation. NOP16 was (F) knocked down, (G) knocked out, and (H) overexpressed in MDA-MB231 cells. Data represents the average ± SEM of the number of mammospheres of two replicates performed in triplicate. *p<0.05, ***p<0.001, ****p<0.0001 as assessed by one-way ANOVA followed by Dunnett’s post-test. **i-k.** NOP16 sustained the migration and invasion ability of breast cancer cells. NOP16 was (I) knocked down, (J) knocked out, and (K) overexpressed in MDA-MB231 cells. (Left) Represent images of migrated and invaded cells stained with crystal violet at 8x. Scale bar, 20 μm. (Right) Quantification of the percentage of cells that migrated or invaded of five fields from each sample performed in triplicate analyzed by Image J. **l**. NOP16 deletion eliminates JMJD3 overexpression-mediated increase of cell proliferation. MDA-MB231 cells were transfected with siRNA for a negative control (N/C) or NOP16 and either an empty vector (E.V.) expression vector or one encoding HA-tagged JMJD3. This graph is a representative experiment of three independent experiments. ****p<0.0001 as assessed by two-way ANOVA followed by Dunnett’s post-test. **m**. The cell survival of NOP16-depleted cells was not affected by EZH2 inhibition. MDA-MB231 cells were transfected with siRNA for a negative control (N/C) or NOP16. One day after seeding, the EZH2 inhibitor DZNep was added to each well for indicated concentration. After 72 hrs, cell viability was measured by WST-1 assay. This graph is a representative experiment of three independent experiments. *p<0.05, ***p<0.001 as assessed by two-way ANOVA followed by Dunnett’s post-test.

**Figure 5.**
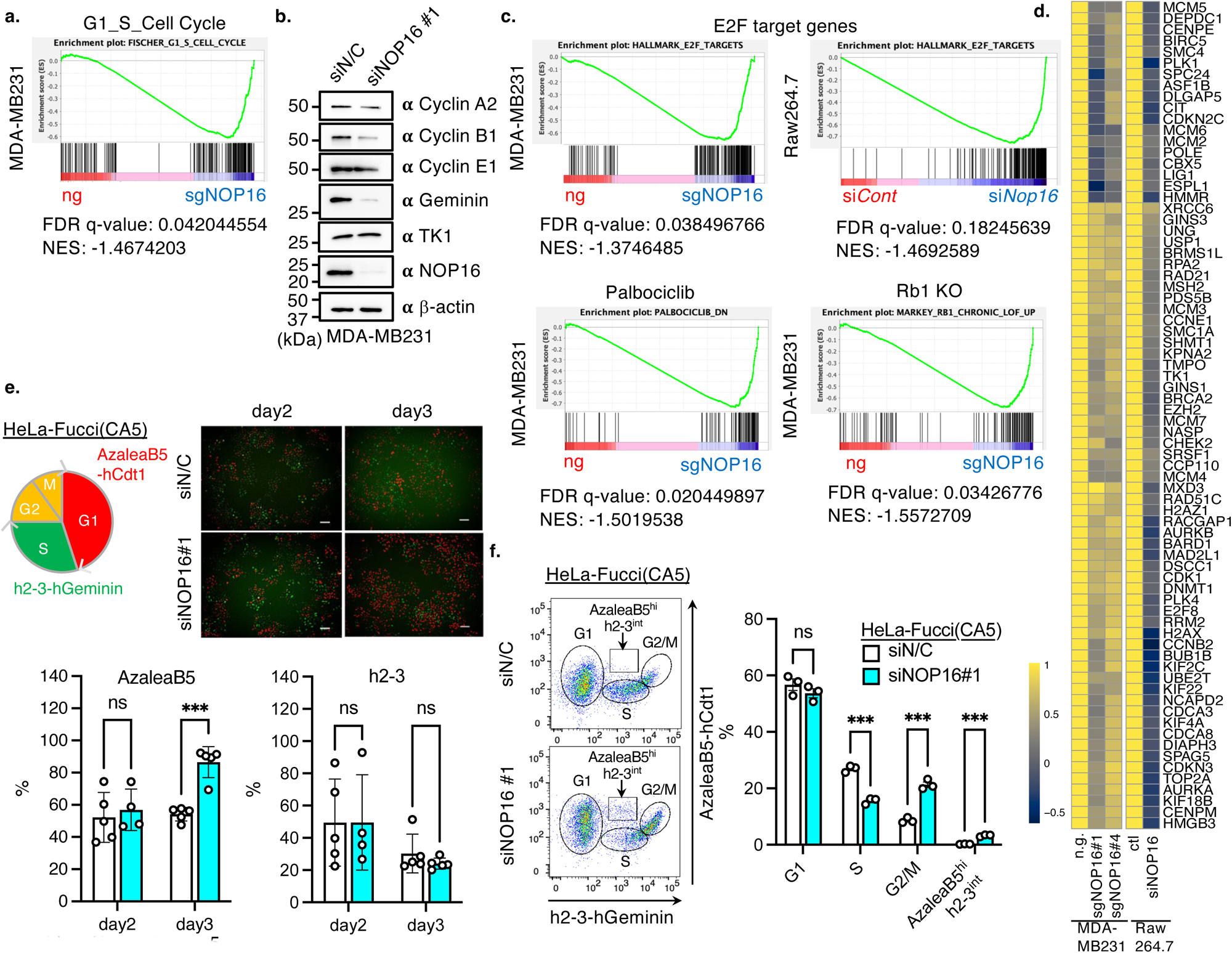
Nop16 regulates expression of cell cycle and E2F target genes. **a.** GSEA analysis shows that NOP16 knockout down regulated the expression of genes categorized as G1-S cell cycle genes. **b.** NOP16 knockdown decreases the protein expression levels of Geminin and Cyclin B1. MDA-MB231 cells were transfected with negative control (N/C) or NOP16 siRNAs (10 pmol) and the protein expression levels of cell cycle regulators by western blots. **c**. GSEA analysis shows that NOP16 depletion in MDA-MB231 and Raw264.7 cells downregulated E2F-target genes (top panels) as well as caused downregulation of genes regulated by RB and CDK4/6, which are upstream of E2F (bottom panels). **d**. A heat map reveals decrease in E2F target genes in response to NOP16 depletion in MDA-MB231 and Raw264.7 cells. **e** and **e**. NOP16 deletion decreases the number of cells in S phase and causes an accumulation of cells in the G2/M phase in HeLa-Fucci(CA5) cells assessed by fluorescent image (**e**) and flow cytometry (**f**). **e**. upper panel displays representative images (scale bar; 20 μm) and lower graph represents percentage of cells at each stage. **f**. left panel displays representative plots graph on right displays percentage of cells in each cell cycle phase. Each column represents the average ± SEM of three independent experiments. ***p<0.001 as assessed by multiple t-test.

**Figure 6.**
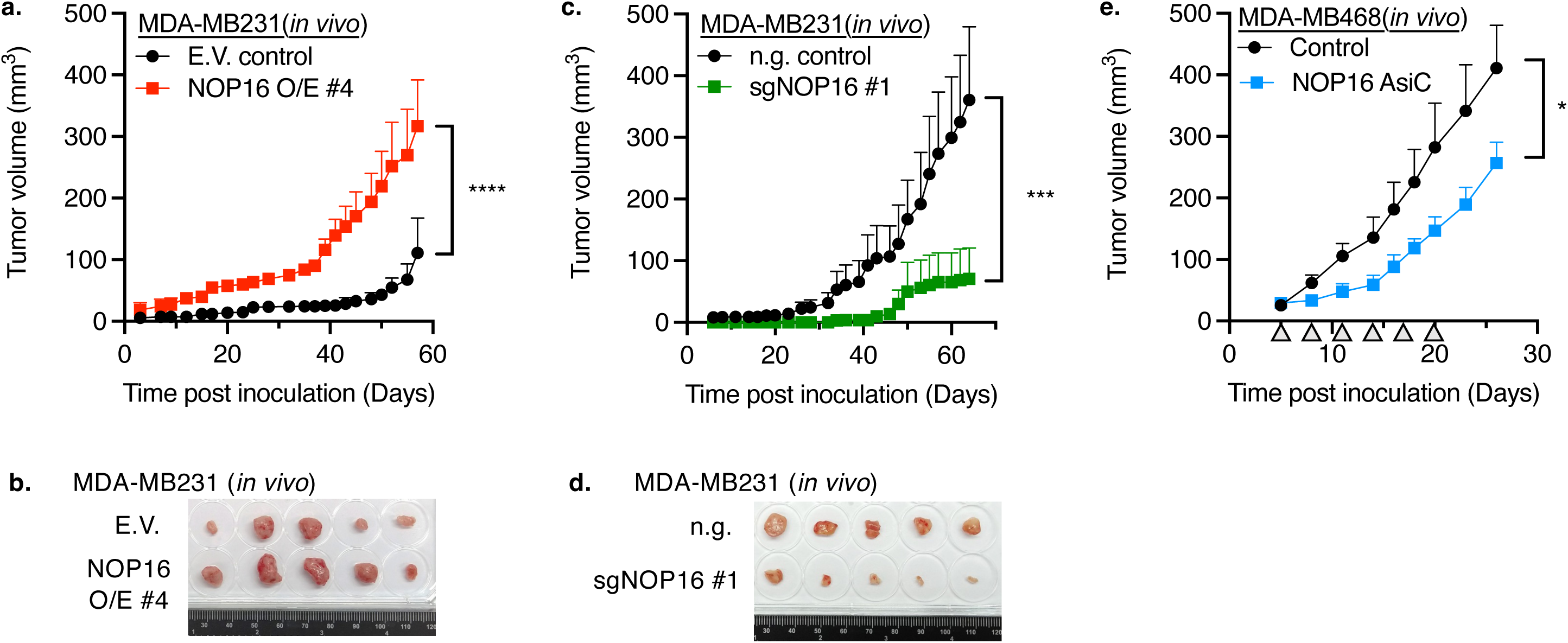
NOP16 is important for breast cancer tumorigenesis *in vivo*. **a** and **b**. Growth of MDA-MB231 tumors in nude mice is increased in response to NOP16 overexpression. MDA-MB231 empty vector (E.V.) control and NOP16 stably overexpressing 3xFLAG-NOP16 cells (NOP16 O/E #4) cells (5×10^6^) were injected in the mammary fat pad of NU/J mice (n=5 per group). A) The size of tumor was monitored every 2-3 days. Data shown as average ± SEM. ****p<0.0001 as assessed by two-way ANOVA followed by Šidák’s multiple comparisons test. B) Individual images of tumor tissues harvested from MDA-MB231 E.V. or NOP16-stable O/E#4-bearing NU/J mice. **c** and **d.** Growth of MDA-MB231 tumors in nude mice is decreased in response to NOP16 deletion. MDA-MB231 no guide (n.g.) control and NOP16 sgRNA#1 (sgNOP16 #1) cells (5×10^6^) were inoculated into the mammary fat pad of NU/J mice (n=5 per group). C) The tumor volume was measured every 2-3 days. Data shown as average ± SEM. ***p<0.001 as assessed by two-way ANOVA followed by Šidák’s multiple comparisons test. D) Individual images of tumor tissues excised from MDA-MB231 n.g. or sgNOP16 #1-bearing NU/J mice. **e.** Palpable tumors generated by MDA-MB468 injection into nude mice displayed reduced growth upon treatment with NOP16-AsiC. MDA-MB468 cells (5×10^6^) were injected in the mammary fat pad of NU/J mice (n=10). Mice were subcutaneously injected with aptamer of control or NOP16-AsiCs (5 mg/kg) in the scruff of the neck every 3 days (indicated as a triangle in figure). The volume of tumor was monitored every 2-3 days. Data shown as average ± SEM. *p<0.05 as assessed by two-way ANOVA followed by Šidák’s multiple comparisons test.

### NOP16 maintains cell proliferation and migration of breast cancer cells

Large scale changes in chromatin modification frequently occur in cancer, including breast cancer ^23^. In particular, H3K27me3, which is associated with gene silencing ^24^, is reduced in breast cancer ^25^. Lower H3K27me3 correlates with poor prognosis ^25^, relapse ^26^, and drug resistance ^27–29^. However, the mechanisms responsible for reduced H3K27me3 and whether and how it contributes to breast tumorigenesis are unknown ^30^. To explore a possible role of NOP16 in breast cancer, we analyzed the TCGA database to determine whether NOP16 expression, which reduces H3K27me3, affects relapse-free survival. Breast cancer patients with lower NOP16 had a significantly better outcome than those with higher NOP16 expression (Fig. 4a). Higher NOP16 expression was associated with the basal subtype of triple negative breast cancer, which has the worst prognosis (Fig. 4b). Genomic alterations, including mutations, structural variants or copy number alterations in cancer-related genes, including *TP53* and *MYC,* also increased with NOP16 expression (Extended Data Fig. 7a).

To define how NOP16 regulates tumorigenesis, we first examined whether NOP16 regulates proliferation, migration and invasivity of breast cancer cell lines. Knockdown or knockout of *NOP16* in MDA-MB231, MCF7, or MDA-MB468 cells significantly decreased cell proliferation (Figs. 4c-d and Extended Data Fig. 7b-h). Changes in proliferation was not restricted to breast cancer cells as knockdown of Nop16 in mouse monocyte macrophage cell line Raw264.7 also caused a decrease in cell proliferation (Extended Data Fig. 7b and e). Conversely, *NOP16* overexpression in MDA-MB231 cells subtly, but significantly, increased cell proliferation (Fig. 4e). The effect of overexpression may have been small because of endogenous NOP16. Knockdown or knockout of NOP16 significantly disrupted mammosphere formation, a measure of tumor-initiating cells (also known as cancer stem cells) (Figs. 4f and 4g). Thus, NOP16 promotes breast cancer cell proliferation and maintenance of the malignant subpopulation of tumor-initiating cells. The effect of NOP16 on tumor cell malignancy was assessed by measuring cell migration across a membrane in response to serum and invasion through Matrigel in Transwell assays. *NOP16* knockdown or knockout decreased both the number of migrating and invading MDA-MB231 cells (Figs. 4i and 4j). Knockout was more efficient than knockdown of *NOP16* (Figs. 4a and 4b) and could therefore explain why the migration phenotype was not observed with one siRNA (Fig. 4i). Conversely, NOP16 stable overexpression slightly increased both the migration and invasion of MDA-MB231 cells (Fig. 4k). Thus, NOP16 promotes malignant properties (cell proliferation, sphere formation, migration and invasion) of breast cancer cells.

### Nop16-mediated increase in proliferation and invasivity depends on H3K27me3 modifiers

Since NOP16 bound to PRC2 and JMJD3, we next investigated whether PRC2 and JMJD3 were involved in NOP16-mediated proliferation. *JMJD3* expression correlates with tumorigenesis and poor prognosis for multiple cancers ^31, 32^, while *EZH2* has oncogenic or tumor suppressive roles depending on context ^33^. However, knockdown of *EZH2* decreases breast cancer cell proliferation ^34, 35^. As expected, overexpression of *JMJD3* promoted proliferation and *NOP16* knockdown decreased proliferation of MDA-MB231 cells. However, *JMJD3* overexpression failed to increase MDA-MB231 proliferation when *NOP16* was knocked down (Fig. 4l), suggesting that the effect of JMJD3 on proliferation depended on NOP16. Similarly, although inhibition of EZH2 reduced MDA-MB231 cell survival, this effect was strongly attenuated when NOP16 was knocked down (Fig. 4m), suggesting that the anti-proliferative effect of EZH2 in this cell line strongly depends on NOP16. Thus, the effect of both EZH2 and JMJD3 on cell proliferation require NOP16.

### Nop16 regulates expression of cell cycle genes and E2F target genes

The more malignant phenotype in NOP16-deficient breast cancer cells correlated with the transcriptome analysis which revealed that *NOP16* knockout down-regulated G1-S cell cycle genes (Fig. 5a and Extended Data Fig. 7i). Decreases in gene expression of cell cycle genes was coupled with decreased protein expression of cell cycle genes, including Geminin, which inhibits DNA replication and is absent during G1 phase and is up-regulated in S phase, and several Cyclin proteins including Cyclin B1, which determines whether cells commit to mitosis (Fig. 5b). Target genes of E2F transcription factors, which regulate the G1->S transition were consistently suppressed in NOP16 knock out in both human and mouse cells (Fig. 5c and 5d). In addition, NOP16 knockout also significantly suppressed the expression of genes regulated by RB and CDK4/6, which are upstream of E2F (Fig. 5c). To measure the direct effect of NOP16 depletion on the cell cycle, we knocked down *NOP16* in HeLa-Fucci (CA5) cells ^36^, which contain a fluorescent cell cycle dependent reporter system. NOP16 depletion decreased the number of cells in S phase and correspondingly increased G2/M phase cells (Fig. 5e, f, and Extended Data Fig. 7j), suggesting that NOP16 deletion affected G1/S transition and mitosis leading to suppression of cell growth.

### NOP16 promotes breast cancer tumor growth

Next we examined whether NOP16 regulates growth of MDA-MB231 xenografts in mice ^37^. MDA-MB231 stably over-expressing *NOP16* or empty vector (E.V.) control cells were subcutaneously injected into nude mice. *NOP16* over-expression significantly increased tumor growth (Figs. 6a and 6b). Conversely, *NOP16* knockout markedly reduced MDA-MB231 tumor growth (Figs. 6c and 6d). To explore whether NOP16 could be a novel therapeutic target for breast cancer, we took advantage of a method of epithelial tumor *in vivo* gene knockdown that links a short 19 nucleotide aptamer that recognizes the epithelial tumor antigen EpCAM on epithelial breast cancers to an siRNA. Subcutaneous injection of EpCAM aptamer-siRNAs (called AsiCs) leads to selective tumor knockdown *in vivo* in EpCAM+ tumors without any apparent toxicity or off-target immune activation ^38–40^. Because MDA-MB231 is mesenchymal and EpCAM-, we used the EpCAM+ epithelial triple negative breast cancer cell line MDA-MB468 to evaluate *in vivo* gene knockdown. EpCAM-AsiCs targeting NOP16 efficiently and selectively knocked down NOP16 in MDA-MB468 *in vitro* (Extended Data Fig. 7k). NOP16 EpCAM-AsiCs also inhibited MDA-MB468 proliferation in a dose dependent manner that mirrored gene knockdown (Extended Data Fig. 7l). Mice bearing MDA-MB468 tumors implanted orthotopically in a mammary fat pad were treated with 5 mg/kg NOP16 EpCAM-AsiCs every three days beginning when tumors became palpable. AsiCs reduced NOP16 mRNA by ∼60% in the tumor *in vivo* as assessed by qRT-PCR (Extended Data Fig. 7m) and significantly suppressed MDA-MB468 tumor growth (Fig. 5e). Thus, NOP16 promotes tumor growth in mice and could be considered as a potential drug target.

## Discussion

Here, we identified NOP16, which contains a 9 amino acid sequence with high homology to the histone H3 tail around Lys27, as a histone H3K27 mimetic. NOP16 binds to H3K27me3 binding protein EED and to the H3K27me3 demethylase JMJD3. NOP16 is itself methylated and methylated lysines are important for NOP16’s interaction with EED. Overexpression of NOP16 decreased global H3K27me3 and conversely NOP16-knockdown or knockout increased H3K27me3 levels and derepressed the expression of genes important for mitosis, DNA replication, cell cycle, cell proliferation, cell division, and apoptosis. In particular, target genes of E2F transcription activators, the master regulators of cell cycle progression, were highly enriched in NOP16-regulated genes. Importantly, dysregulation of H3K27me3 went hand in hand with cellular phenotypes important for breast cancer growth and malignancy. Moreover, NOP16 knockdown using tumor-directed EpCAM-AsiCs attenuated tumor growth *in vivo* in triple negative breast cancer orthografts in mice. Together these findings identified an endogenous histone mimetic that regulates H3K27me3, cell cycle progression and tumor growth.

Biochemical experiments revealed that NOP16 interacts with the PRC2 complex through the WD40 domain of EED (Fig. 1e), and that Lys 29, 106, and 107 of NOP16 enhance this binding (Fig. 2c). The WD40 domain of EED is essential for binding to H3K27me3 and recruiting the PRC2 complex to specific genomic regions to propagate, and/or maintain H3K27me3 ^18^. Synthetic NOP16 peptides that were di or tri-methylated at Lys29 (NOP16 K29 me2 or me3) directly bound to the WD40 domain of EED (Fig. 2e). However, mass spectrometry and western blotting analysis failed to detect methylation of NOP16 at Lys29. It is likely that this residue is methylated, but methylation was not detected. Mass spectrometry is inherently non-exhaustive and doesn’t detect all possible peptides and the methyl specific antibody could have preference for some methylated lysines over another (for instance mono or di-methylated lysines rather than trimethylated lysines). Mass spectrometry and *ex vivo* experiments, however, also indicated that NOP16 Lys 106 and 107 were also methylated (Fig. 2b) and suggested that NOP16 and EED interact in other regions besides the homologous sequence near the N-terminus of NOP16. Di- and trimethylated lysine 29 NOP16 peptides bound to EED, and substitution of all three lysine residues (K29, K106 and K107) with alanine was necessary to abrogate NOP16 binding to EED. These findings suggest that K29 of NOP16 is indeed methylated. However, future structural studies and identification of the enzymes that methylate these NOP16 residues are needed to clarify NOP16’s interaction with EED, how it is regulated, whether K29 is methylated and its role in NOP16 binding and function.

NOP16 regulated the level and distribution of H3K27me3 (Fig. 3). The simplest model is that NOP16 competes with H3K27me3 for EED binding, which would inhibit the maintenance or propagation of H3K27me3. NOP16 could also affect the enzymatic activity of PRC2 or JMJD3. EED recognizes tri-methylated lysine within an Ala-Arg-Lys-Ser (ARKS) motif and thus binds not only to H3K27me3 but also to H3K9me3 and H1K26me3 ^41^. H3K27me3 peptides have been shown to enhance, while H1K26me3 peptides repress, PRC2 methyltransferase activity ^41^. NOP16 binding to EED could affect PRC2 methyltransferase activity. Alternatively, NOP16 could regulate H3K27me3 levels through its interaction with the H3K27me3 demethylase JMJD3. Because NOP16 bound to JMJD3’s JmjC catalytic domain, NOP16 might also regulate its enzymatic activity and/or localization. NOP16 depletion affected cell cycle genes generally and more specifically E2F target genes. The molecular basis for this possible selectivity for cell cycle regulating genes is unclear. JMJD3 was previously shown to decrease H3K27me3 and specifically regulate genes within the pRB-E2F pathway ^42^. Therefore, NOP16 could regulate cell cycle genes by disrupting JMJD3’s capacity to demethylate H3K27me3 at E2F target genes.

Genes involved in lipid metabolism were highly enriched in NOP16-regulated genes. A recent report showed that NOP16 associated with ATP-citrate lyase (ACLY) and altered lipid metabolism in cancer-associated fibroblasts (CAFs) in a colorectal cancer model of liver metastasis ^22^. This work suggested that NOP16 upregulated acetyl-CoA and increased H3K27 acetylation indirectly^22^. This potential NOP16 activity could complement its role in regulating H3K27 methylation to enhance NOP16’s effect on lipid metabolism. Future studies will need to examine whether NOP16 inhibits methylation of H3K27 directly by competing for EED binding and/or indirectly by increasing its acetylation.

Drugs that mimic and displace histone modified peptides or inhibit histone modifying enzymes are approved or being developed as potential therapeutic and immunomodulatory drugs based on their ability to sequester or inhibit the chromatin regulatory machinery ^43^. NOP16, a natural histone H3K27 mimetic, could be a novel therapeutic target for manipulating chromatin architecture and gene expression. Consistent with this idea, here we showed that knockout or tumor-targeted knockdown of *NOP16* in a triple negative breast cancer model inhibited tumor growth (Fig. 5). Since partial *NOP16* knockdown (∼60%) significantly attenuated tumor growth, more efficient blockade of NOP16 could be more effective. Current cytotoxic, targeted or immunological treatments of triple negative breast cancer are inadequate and need to be improved ^44^. Combining NOP16 knockdown with existing therapies could improve therapeutic efficacy. NOP16 is a frequent top “hit” in CRISPR screens in multiple cancer cell lines ^16^, further supporting NOP16 as a potential drug target that might be active in multiple types of cancer.

Here we identified NOP16 as a novel endogenous histone mimic, which can regulate H3K27me3 and gene expression. Until now, histone mimetics that disrupt the chromatin regulatory machinery have only been reported to be encoded by viruses - Influenza A ^9^. and recently SARS-CoV2 ORF8 ^45^. Our identification of an endogenous histone mimetic raises the possibility that additional endogenous histone mimetic proteins may exist. NOP16 and other histone mimetics offer the possibility of an endogenous tunable mechanism of regulating chromatin modifications. Manipulating histone mimetics might offer better therapeutic efficacy with less toxicity than inhibiting histone methyltransferases or demethylases since these enzymes play such a critical role in normal cell development and maintaining cellular identity.

## Acknowledgements

This work was supported by JSPS (Overseas Research Fellowships, Grant-in-Aid for Research Activity Start-up: 21K20841, Grant-in-Aid for Transformative Research Areas(A): 22H05603), The Uehara Memorial Foundation (Post-doctoral Fellowship) and Tasaki Memorial Research Grant to K.T. and a DFG research fellowship to D-J. L.. Work by M.M. and S.H.S. was supported in part by Harvard Catalyst | The Harvard Clinical and Translational Science Center (NIH award UL1TR002541). This work was supported by NIH grant DP2AG055947 to E.L.G..

## Author contribution

K.T. and E.L.G conceived and planned the study and wrote the paper. K.T., H.O., J.L. and E.L.G designed experiments. K.T. performed biochemical and cellular functional analysis of NOP16, and prepared CUT&RUN and RNA-seq experiments. D-J. L. performed mouse xenograft experiments and was aided by Y.Z. and Z.L. and supervised by J.L. M.F.T. produced Figs. 1F, 2D, S2B and S2D and was aided by Z.L.L.. The bioinformatic analysis of RNA-seq and CUT&RUN experiments was performed by M.H.R. and M.M. and was supervised by S.H.S.. C.P.D. supervised CUT&RUN-seq and performed NGS sequencing for CUT&RUN-seq. J.N., E.S. and J.A H. helped purify proteins and perform *in vitro* binding assays. G.B.H. performed biochemical analysis supervised by K.T.. S.D. performed comparative analysis of NOP16 depletion in human and mouse cell lines.

## Declaration of interests

The authors declare no competing interests.

## Resource Availability

### Lead contact

Further information and requests for resources and reagents should be directed to and will be fulfilled by the lead contact, Eric Lieberman Greer

### Materials availability

All unique/stable reagents generated in this study are available from the lead contact without restriction.

## Methods

### Cells, Reagents

HeLa-Fucci(CA5) cells were provided by the RIKEN BRC. HEK293T, HEK293FT, MCF7, MDA-MB231, HeLa, HeLa-Fucci(CA5), A549, Raw264.7 cells were cultured in Dulbecco’s modified Eagle’s medium (D-MEM, 11995-065, Thermo Fisher Scientific) high glucose medium with 10 % heat-inactivated fetal bovine serum (FBS) and antibiotics (100 units/ml penicillin and 100 μg/ml of streptomycin) at 37 °C with 5% CO_2_. MDA-MB468 cells were cultured in RPMI1640 (22400105, Thermo Fisher Scientific) supplemented with 10 % heat-inactivated FBS and antibiotics (100 U/mL penicillin G and 100 μg/mL streptomycin), 6 mmol/L HEPES, 1.6 mmol/L l-glutamine, 50 μmol/L β-mercaptoethanol (Sigma-Aldrich) at 37 °C with 5% CO_2_. DZNep was purchased from Sigma-Aldrich.

### Antibodies

mouse anti-FLAG M2 (F1804, Sigma, 1:1000), mouse anti-β actin (A5441, Sigma, 1:5000), rat anti-HA-Peroxidase (3F10, Sigma, 1:2000), rabbit anti-HA antibody (H6908, Sigma, 1:1000), rabbit anti-Pan methyl Lysine antibody (ab7315, Abcam, 1:1000), mouse anti-NOP16 (ABIN526754, Abnova, 1:1000), rabbit anti-NOP16 (H00051491-D01P, Abnova, 1:1000), mouse anti-EZH2 (AC22, Active Motif), rabbit anti-EZH2 (D2C9, Cell Signaling Technology, 1:1000), anti-EED (05-1320, AA19, Sigma-Aldrich, 1:1000), anti-JMJD3 (ab38113, Abcam, 1:1000), anti-H3K4me3 (ab8580, Abcam, 1:1000), anti-H3K9me3 (ab8895, Abcam, 1:1000), rabbit anti-H3K27me3 (07-449, EMD Millipore, 1:1000), rabbit anti-H3K27Ac (ab4729, Abcam, 1:1000), anti-H3K36me3 (ab9050, Abcam, 1:1000), rabbit anti-H3 (ab1791, Abcam, 1:5000), anti-SP1(#9389, D4C3, Cell Signaling Technology, 1:1000), anti-HSP90 (#4874, Cell Signaling Technology, 1:1000), anti-Geminin (#52508, Cell Signaling Technology, 1:1000), anti-Thymidine Kinase 1(TK1, #28755, Cell Signaling Technology, 1:1000), anti-Cyclin A2 (#91500, Cell Signaling Technology, 1:1000), anti-Cyclin B1 (#12231, Cell Signaling Technology, 1:1000), anti-Cyclin E1(#20808, Cell Signaling Technology, 1:1000). For flow cytometry, PE-conjugated anti-human CD24 (311105, ML5, BioLegend, 1:200), FITC-conjugated anti-mouse/human CD44(103021, BioLegend, 1:200)

### RNA interference (RNAi)

Knockdown of NOP16 was carried out using small interfering RNA (siRNA). Each siRNAs (siNOP16#1: s28205, siNOP16#2: s28206, si*Nop16* for Mus: s203208, siN/C: Silencer Select Negative Control #1) were purchased from Invitrogen. Following the manufacturer’s instructions, cells were transfected with siRNAs (for siNOP16: 10 nM) using Lipofectamine 3000 or Lipofectamine RNAi Max reagent.

### Aptamer–siRNA chimeras (AsiC)

The long strand of the AsiC synthesized with 2′-fluoropyrimidines was annealed to the short antisense strand (TriLink Biotechnologies) with a 2-fold molar excess of the short strand. After the long strand was heated to 95°C for 10 min, the short strand was mixed with the long stand to anneal at 65°C for 7 min. The mixture was cooled down at room temperature for 20 min. The annealed AsiC duplexes were purified using Illustra MicroSpin G-25 columns (GE Healthcare Life Sciences). NOP16 EpCAM-AsiC sense and antisense sequence were as follows; NOP16-EpCAM sense: GCGACUGGUUACCCGGUCG **UUU** GGACCUCAUUGACUAUGUAdTdT, NOP16 antisense: UACAUAGUCAAUGAGGUCC dTdT. Bolded UUU is linker sequence.

### Gene knockdown using AsiC

To measure *in vitro* NOP16 EpCAM-AsiC-mediated gene silencing, MDA-MB468 cells (1×10^4^ cells/well) were incubated with NOP16 EpCAM-AsiC (1 - 4 μmol/L) or EpCAM aptamer (4 μmol/L). Seventy-two hours after treatment, gene knockdown was assessed by measuring mRNA expression using RT-qPCR, and cell viability was measured by CellTiter-Glo assay (Promega). To assess *in vivo* gene silencing, tumors were collected from mice treated with NOP16 EpCAM-AsiC or EpCAM aptamer. RNA was extracted from each tumor and gene knockdown efficiency was evaluated by RT-qPCR.

### Establishment of NOP16-deficient cells and overexpression cell lines

HEK293T, MCF7 and MDA-MB231 *NOP16*-deficient cell line were generated by CRISPR/Cas9 system using gRNA targeting NOP16 exon 1 (sgRNA1; 5’-GGTGGTCAGCGCGATGCCCAAGG-3’, sgRNA4; 5’-GAACCGGAATGCTCGACGG-3’). Oligonucleotide duplex corresponding to these gRNAs were inserted into the pSpCas9n(BB)-2A-Puro (PX459) V2.0 vector (62988, Addgene) or lentiCRISPR v2 vector (52961, Addgene). For gene knockout in HEK293T cells, 1 μg of pX459-NOP16 sgRNA was transfected into cells using Lipofectamine 3000 (Thermo Fisher Scientific). Twenty-four hours after transfection, we selected the positive cells using 1 μg/ml Puromycin for three days. For gene knockout in MDA-MB231 cells, lentivirus encoding Cas9 and sgRNA were produced. 1 μg of lentiCRISPR v2-NOP16 each sgRNA, packaging vectors (1 ug total): VSV-G (0.25 μg), HgDM2 (0.25 μg), tat (0.25 μg), CMVRaII (0.25 μg) were transfected into HEK293T cells. Cell supernatant containing lentivirus was collected at 48-72 hr after transfection and filtered with 0.45□m filter (Millipore). For infection into MDA-MB231 cells, cells were seeded at 2×10^5^ cells per 6-well plate, and added 1 ml viral supernatant plus 1 ml D-MEM containing 8 μg/ml of Polybrene. Forty eight hours after infection, transduced cells were selected in D-MEM containing 1 μg/ml Puromycine for 2 days. For establishment of stable cell lines overexpressing, MDA-MB231 cells were transfected with 1 μg p3xFLAG-CMV10-NOP16 full-length or p3xFLAG-CMV10-empty vector, and each line were selected and expanded in D-MEM containing 1 μg/ml G418 selection for two weeks. The levels of protein expression were confirmed by western blot.

### Histone preparation

Histones were purified using acid extraction ^46^. Cells (1-5×10^6^) were lysed in hypotonic lysis buffer (10 mM Tris–HCl pH 8.0, 1 mM KCl, 1.5 mM MgCl_2_, 1 mM DTT and protease inhibitors (Roche)) for 30 min on rotator at 4 °C. Nuclear pellet was collected by centrifuge at 10,000 rpm for 10 min at 4 °C, and then re-suspended and incubated with 0.2 M H_2_SO_4_ for overnight on rotator at 4 °C. After removing nuclear debris by centrifuge at 14,000 rpm for 15 min at 4 °C, the supernatant containing histones were added TCA (final concentration of TCA is 33%), and incubated for overnight at 4 °C. Precipitated histone was collected by centrifuge at 14,000 rpm for 15 min at 4 °C, and washed with ice-cold acetone twice. After air dry, histone pellets were dissolved in ddH_2_O, and protein concentration was determined by Bradford assay.

### Immunoprecipitation and western blotting

Cells were lysed in lysis/wash buffer (20 mM Tris-HCl pH 7.6, 125 mM NaCl, 1 mM EDTA, 10% glycerol, 1% NP-40, 1 mM PMSF and protease inhibitors) on ice for 30 min, and then centrifuged at 14,000 rpm for 20 min at 4 °C. A part of lysate were stored for “input”. Antibody (1:100) was added to cell lysate, and the lysate was incubated with rotation for 2 hours at 4°C. Pre-washed Dynabeads Protein G (Invitrogen) or Protein G Sepharose (Cytiva) were added into the lysate with antibody and incubated with rotation for overnight at 4 °C. And then, beads were washed 4 times with lysis/wash buffer. For elution, samples were boiled in lysis/wash buffer added with 2x SDS sample buffer (125 mM Tris-HCl pH 6.8, 35% glycerol, 4% SDS, BPB) containing 2-mercaptoethanol for 5 min at 98°C and the supernatant was collected as “IP sample”. Same amounts of samples were subjected to 6-15 % SDS-PAGE, and transferred to nitrocellulose or PVDF membrane. For detection of histone protein, Ponceau-S (Beacle) staining was performed as loading control. The membrane was blocked with 5% skim milk in TBS (20 mM Tris-HCl pH 7.6, 150 mM NaCl) and then incubated with primary antibody for overnight at 4°C. After 3 times wash with TBS-T (20 mM Tris-HCl pH 7.6, 150 mM NaCl, 0.1% Tween-20), the membrane was incubated with HRP-conjugated secondary antibody (1/10000) for 1 hour at room temperature. After 3 times wash, the membranes were treated with Immobilon ECL Ultra Western HRP Substrate (Millipore) or ECL Prime Western Blotting Detection Reagent (Cytiva). The signal of blots was detected with ChemiDoc Touch (Bio-Rad).

### Protein expression and purification

Full-length human NOP16, EZH2, EED, SUZ12 cDNA sequence was cloned into pET28a or pGEX-4T1 vector. Each deletion or mutant was produced by PCR using Pfu Ultra Ⅱ Fusion HS DNA Polymerase (Agilent Technology 600670). Vectors were transfected into T7 Express lysY Competent E. coli (NEB C3010). Overnight induction of protein expression was carried out with 0.5 mM IPTG at 18°C. Bacteria were harvested at 4000 rpm, 4°C and the cell pellet was resuspended in protein purification lysis buffer (50 mM Tris-HCl pH 7.5, 0.25 M NaCl, 0.1% Triton-X, 1 mM PMSF, 1 mM DTT, and protease inhibitors). The lysate was sonicated 5 times in 30 s on/off cycles and then centrifuged at 12,000 rpm for 20 minutes. Lysates were incubated with glutathione Sepharose 4B beads (Sigma) for GST-tagged protein or Ni-NTA Agarose beads (QIAGEN) for His-tagged protein. Proteins and beads were washed 3 times with protein purification lysis buffer before incubating the beads with elution buffer (12 mg/ml Glutathione in protein purification lysis buffer, pH 8.0 for GST-tagged protein, 20mM Imidazole in protein purification lysis buffer for His-tagged protein) for 30 minutes. Eluates were dialyzed overnight at 4°C with enzyme storage buffer (50 mM Tris-HCl pH 7.6, 100 mM KCl, 5 mM MgCl_2_, 0.1% NP-40, 1 mM PMSF, 1 mM DTT, 10 % glycerol) and were subsequently stored at -80°C. Bradford assays and SDS-PAGE followed by Imperial staining (Thermo Scientific #24615) was performed to determine integrity and quantity of purified proteins.

### GST-Pull down assay

Glutathione-Sepharose 4B Beads were blocked with 3.5% BSA at 4 °C for 1 hour. 5 μg of GST-tagged protein and 5 μg of His-tagged protein were added with 1xTAP Wash Buffer (50 mM Tris (pH 8.0), 100 mM KCl, 10% glycerol, 5 mM MgCl_2_, 0.2 mM EDTA, 0.1% NP-40) and rotated at 4 °C for 1 hour. Beads were washed 4 times with cold 1xTAP Wash Buffer and then 1 time with cold 1xPBS. Samples were eluted in 1x SDS sample buffer, boiled, loaded on an SDS-page gel, and Imperial stained.

### Peptide binding assay

Pierce Streptavidin Magnetic Beads (Thermo Scientific 88817) were washed 3 times in reaction buffer (50 mM Tris-HCl pH 7.5, 150 mM NaCl, 0.2 mM EDTA, 0.1% NP-40). Each 1 μg of biotinylated peptide (synthesized by Thermofisher) and 10 μg of bacterially purified GST tagged protein were added and allowed to incubate at 4°C for 2 hours. Beads were washed 4 times with reaction buffer. Samples were eluted in 1x SDS sample buffer, boiled, loaded on an SDS-page gel, and Imperial stained.

### Microscale thermophoresis

Binding affinities were estimated by microscale thermophoresis using a Monolith NT.115 device (NanoTemper, Germany). We measured the thermophoretic movements of labeled molecules within a temperature gradient, inside standard capillaries (NanoTemper, MO-K002). EED and JMJD3 proteins were labeled with Protein Labeling kit RED-NHS (NanoTemper, MO-L001). At least three independent experiments were performed for each binding reaction.

### RT-qPCR and RNA sequencing

Total RNA was extracted using the Direct-zol RNA Miniprep kits (Zymo Research) following the manufacturer’s instruction. The cDNA was synthesized by reverse transcription of 400 ng of RNA using the High Capacity cDNA Transcription Kit (ABI) with random primers. qPCR was performed using the iTaq Universal SYBR (Bio-Rad), power SYBR Green PCR Master Mix (Life Technologies) or SsoFast EvaGreen Supermix (Bio-Rad) and the CFX96 Touch Real-Time PCR Detection system (Bio-Rad), Step-one Real-time PCR system (Life Technologies) or Bio-Rad C1000 Thermal Cycler (Bio-Rad). The sequence of primers qPCR were as follows; NOP16 qF: ATG GAG GTG GAC ATA GAG GAG AGG, NOP16 qR: TCA ATG AGG TCC CGA GAC AGA GTA T, GAPDH qF: ACC ACA GTC CAT GCC ATC AC, GAPDH qR: TCC ACC ACC CTG TTG CTG TA. The RNA expression levels were normalized to GAPDH. Total RNA for RNA sequencing was sent for polyadenylate selection, library preparation, and sequencing on Illumina NovasSeq 6000 platforms at Novogene.

### Read mapping and expression level estimation for RNA-seq

All samples were processed using an RNA-seq pipeline implemented in the bcbio-nextgen project (https://bcbio-nextgen.readthedocs.org/en/latest/). Raw reads were examined for quality issues using FastQC (http://www.bioinformatics.babraham.ac.uk/projects/fastqc/) to ensure library generation and sequencing data were suitable for further analysis. If necessary, adapter sequences, other contaminant sequences such as polyA tails and low quality sequences were trimmed from reads using cutadapt ^47^. Trimmed reads were aligned to the hg38 build of the human genome using STAR ^48^. Alignment quality was checked for evenness of coverage, rRNA content, genomic context of alignments (for example, alignments in known transcripts and introns), complexity and other quality checks. Expression was quantified with Salmon ^49^ to identify transcript-level abundance estimates and then collapsed down to the gene-level using the R Bioconductor package tximport ^50^. Principal components analysis (PCA) and hierarchical clustering methods validated clustering of samples from the different NOP16 manipulations and controls.

### Differential gene expression and functional enrichment analysis for RNA-seq

Differential expression was performed at the gene level using the R Bioconductor package DESeq2^51^. For each comparison significant genes were identified using an FDR threshold of 0.05. Heatmaps of significant genes were plotted using the pheatmap package in R. Functional enrichment was evaluated using the clusterProfiler package in R ^52^. Gene Ontology (GO) over-representation analysis was performed to identify relevant biological processes. Gene set enrichment analysis (GSEA) was used to identify coordinated changes of gene expression in pathways.

### Cleavage Under Targets and Release Using Nuclease (CUT&RUN)

Cells were washed with cold 1xPBS and lysed with nuclear extraction buffer (20 mM HEPES pH 7.9, 10 mM KCl, 0.1% Triton X-100, 3 mM MgCl_2_, 0.5 mM Spermidine, 10mM Sodium Butyrate and protease inhibitors) with rotation at 4°C for 10 min. Nuclei were collected by centrifuge at 2000 rpm for 10 min at 4 °C and counted. 1.5×10^5^ nuclei were stored in cold wash buffer (20 mM HEPES pH 7.5, 150 mM NaCl, 0.2% Tween-20, 0.1% BSA, 0.5 mM Spermidine, 10mM Sodium Butyrate and protease inhibitors) in 1.5 mL low-binding tube (Eppendorf, 022-43-102-1). Nuclei were incubated with concanavalin A beads (Bangs Laboratories) in binding buffer (20 mM HEPES pH 7.9, 10 mM KCl, 1 mM CaCl_2_, 1 mM MnCl_2_) at room temperature for 10 min. After removing the supernatant on the magnetic stand, the beads were added with each diluted antibody (1:100) in 50 μl cold antibody buffer (wash buffer supplemented with 0.1 % Triton X-100 and 2 mM EDTA) per sample and incubated at 4 °C for overnight. Beads were washed with antibody buffer and incubated with 700 ng / mL pA-MNase at 4 °C for 1 hour. After 2 washes with Triton-wash buffer (wash buffer supplemented with 0.1 % Triton X-100), CaCl_2_ was added to a final concentration of 3 mM to activate pA-MNase and the reaction was carried out by incubating on a 0 °C cold metal block for 30 min, and then stopped by adding 2xSTOP buffer (340 mM NaCl, 20 mM EDTA, 4 mM EGTA, 0.04 % Triton X-100, 20 pg/mL Yeast-Spike-in DNA, 100 μg/ml RNase A). DNA-protein complexes were released by incubation at 37°C for 20 min and then beads were captured on the magnetic stand, and the supernatant was transferred to fresh 1.5 mL low-binding tube. Samples were added with 1/100 vol. of 10 % SDS and 1/100 vol. of 20 mg/ml Proteinase K and shaken at 65°C for 1 hour in thermomixer at 500 rpm. DNA were extracted with Phenol-Chloroform and purified by ethanol precipitation. Briefly, for library preparation, end repair was conducted for 15 min at 12°C and 15 min at 37°C, followed by dA-tailing for 20 min at 72°C. After adaptor ligation using TruSeq DNA single indexes (Illumina) for 15 min at 20°C, adapter-ligated DNA was cleaned up by Ampure XP beads (Beckman Coulter). Libraries were PCR amplified for 14 cycle using KAPA HS HIFI polymerase. After cleaning up of PCR product by Ampure XP beads, DNA amount of library was measured by Qubit and size distribution of DNA was determined by Agilent TapeStation. The samples were put into a pooled library (0.5 nM) and sequenced on Nextseq 500 platform (Illumina) with paired-end 75-base pair reads.

### Read mapping and peak calling for CUT&RUN

CUT&RUN-seq data quality was evaluated using FASTQC (http://www.bioinformatics.babraham.ac.uk/projects/fastqc/), and if required reads were trimmed with Atropos ^53^. High quality sequencing reads were mapped to the hg38 human genome build using Bowtie2 v2.4.1 ^54^. Alignments were filtered to retain only reads with a unique mapping to the genome and peak artifacts were filtered out using the ENCODE blacklist regions ^55^. Peaks were called with MACS2 v2.2.7.1 ^56^ using the broad peak calling sub-command and peak cutoff was set to q < 0.05. To assess the reproducibility of peaks across replicates, two methods were implemented. First, we plotted the peak rank versus the signal enrichment to identify the signal strength of each replicate against each other. Second, we computed overlaps between replicates using a liberal threshold of 1bp overlap.

### Peak annotation and visualization for CUT&RUN

Bigwig tracks were created for each individual file using the deepTools suite ^57^ with CPM normalization. Peaks annotation and visualization were performed using ChIPseeker ^58^. Annotation was generated based on proximity to neighboring genes’ transcription start sites (TSS). A custom transcript database (TxDb) was created using a GTF file obtained to match the version of the hg38 genome used during alignment. Consensus peaksets for each group were defined as regions overlapping across all replicates by a minimum of 1 bp. Visualizations were generated for each consensus set to identify changes between groups.

### Differential enrichment analysis for CUT&RUN

The R Bioconductor package DiffBind ^59^ was used to identify binding regions that show a significant change in the H3K27me3 mark between NOP16 manipulation samples and respective controls. Default normalization was applied and the DESeq2 was used to make all comparisons. Significant regions were identified at FDR < 0.05

### Cell proliferation assay

For knockdown condition, 5×10^5^ Cells were transfected with each siRNA using Lipofectamine 3000 or Lipofectamine RNAiMax. After 24 hours from transfection, 5×10^4^ cells per well were plated on a 6 well plate. For NOP16-stable expression line or NOP16 sgRNA-mediated bulk knock out line, 5×10^5^ Cells were per well were plated on a 6 well plate. After 24 hours, cells were trypsinized and the cell number was counted. 5×10^4^ cells per well were plated on a 6 well plate. This cell number was set as day 0. On the indicated days, the cells were washed with PBS, trypsinized, stained with trypan-blue, and counted. For DZNep treatment following gene knockdown, 1000 cells per well were plated on a 96 well plate. On next day after seeding, DZNep were added to cell supernatant for indicated final concentration. Three days after DZNep addition, cell viability was measured by Premix WST-1 Cell proliferation Assay System.

### Migration / Invasion assay

For migration assay, Falcon Permeable Support for 24 Well Plate with 8.0 μm Transparent Polyester (PET) Membrane (Corning #353097) was used as according to the manufacturer’s instructions. 1×10^5^ Cells in 200 μl serum-free DMEM were seeded on the upper wells, and 700 μl DMEM supplemented 10% FBS was placed on the lower wells. For invasion assay, BioCoat Matrigel Invasion Chambers with 8.0 μm Pore Polyester (PET) Membrane (Corning #354480) were used following the manufacturer’s instructions. 1×10^5^ Cells in 500 μl serum-free DMEM were seeded on the upper wells, and 750 μl DMEM supplemented 10% FBS was placed on the lower wells. After incubation for 16 hr, cells were fixed in 4% PFA for 10 min and stained with 1 % Crystal violet (Sigma-Aldrich V5265) for 30 min at room temperature. Non-migrating cells on the upper surface of the filter were removed with cotton swabs. Cells were visualized at x8 magnification using stereo Discovery V8 microscope (ZEISS), and analyzed by Image J (version 1.53) software.

### Sphere formation assay

MDA-MB231 cells were washed with Hank’s Balanced Salt Buffer and harvested with cell scraper. Cells were re-suspended with MammoCult Human Medium (Stem cell technology #05620), and then 5000 cells per well were seeded on 6 well Ultra-low adherent plate (Corning #3471) and cultured for 1 week. Sphere were gently harvested and counted under microscope.

### Cell cycle analysis with HeLa-Fucci(CA5) cells

HeLa-Fucci(CA5) cells were plated in 6 well plates (5×10^4^ cells/well) and transfected with each siRNA using Lipofectamine RNAiMax. 48 or 72hrs after transfection, fluorescent images were visualized at x4 magnification using BZ-X810 Fluorescence microscope. 72hrs after transfection, cells were also analyzed on FACS Verse flow cytometer (BD Bioscineces) and data analysis was performed using FlowJo v10.6 (BD).

### Mouse studies

All animal experiments were conducted in compliance with all the ethical regulations and were approved by the Harvard Medical School Institutional Animal Care and Use Committee. The mice were housed in the Harvard Center for Comparative Medicine. Female nude NU/J mice (6-8 weeks old) were purchased from the Jackson Laboratories. For orthotopic tumor challenge, NOP16-overexpressing, NOP16-knocked out, or control MDA-MB231 cells (5 × 10^6^ cells/mouse) were injected into the 4th mammary fat pad of mice (N = 5). To determine the longer-term anti-tumor efficacy of NOP16 EpCAM-AsiC, MDA-MB468 cells (5 × 10^6^/mouse) were injected into the 4th mammary fat pad of mice. On day 5 post tumor challenge, mice were treated with the NOP16 EpCAM-AsiC (5 mg/kg) or EpCAM aptamer (5 mg/kg) every third day (N = 5). Tumor growth was monitored by measuring the perpendicular diameters of tumors three times per week.

## Notes

### Competing Interest Statement

The authors have declared no competing interest.

