## Supplemental Figures for "NOP16 is a histone mimetic that regulates Histone H3K27 methylation and gene repression"

**Supplemental information**

**Extended Data Figure 1. NOP16 interacts with H3K27 modifiers**

**a.** Alignment of NOP16 shows conservation throughout the phylum chordata. "*", ":", and "." indicate identical, strong similarity, and weak similarity, respectively. **b.** NOP16 is evenly expressed in different breast cancer and other epithelial cell lines as assessed by western blotting. **c.** NOP16 is present in both the soluble nuclear and chromatin bound fractions. Fractionated cell lysate from MDA-MB231 cells were analyzed by western blotting. **d.** EED binds to both the N and C terminus of NOP16 *in vitro*. **e.** Amino acids 1-22 and 23-44 of NOP16 bind to EED *in vitro.* **f**. NOP16 directly binds to the JmjC domain of JMJD3. Samples from GST pull downs and input were subjected to SDS-PAGE, and blotted with an anti-His antibody.

**Extended Data Figure 2. NOP16 methylation affects binding of EED but not JMJD3**

**a.** Peptide sequence detected by mass spectrometry with methylated lysines highlighted in red. **b.** Di and tri methylated lysine 29 biotinylated peptides of amino acids 25-32 of NOP16 directly bound to GST-tagged EED *in vitro.* **c**. Both methylated and unmethylated biotinylated NOP16 peptides of amino acids 25-32 directly bound to GST-tagged JMJD3 JmjC *in vitro.* 1 μg of synthetic NOP16 peptides (25-32 amino acids) which were unmethylated (me0), mono(me1)-, di(me2)-, or tri(me3)-methylated were incubated with 10 μg of GST-tagged EED or GST-tagged JMJD3 JmjC protein. Binding was detected by imperial staining after streptavidin pull-down or by western blotting with a GST antibody. **d.** Microscale thermophoresis of the Jumanji domain of JMJD3 and methylated or unmethylated NOP16 or H3K27me3 peptides show that JMJD3 JmjC has highest affinity for H3K27me3 than NOP16K29me3 than NOP16K29me0 than NOP16 K29me1than NOP16K29me2. This table represents the mean +/- SD of 5-7 independent experiments.

**Extended Data Figure 3. CUT&RUN analysis of NOP16 manipulated cells**

**a.** Knock down of NOP16 increases H3K27me3 in MDA-MB231 cells. MDA-MB231 cells were transfected with negative control (N/C) or NOP16 siRNAs (10 pmol) and levels of H3K27me3 was determined by western blotting of extracted histones. Ponceau-S staining is used to display total histone levels while NOP16, EZH2 and beta-actin control expression is displayed by western blotting of whole cell lysates (WCL). **b.** NOP16 overexpression caused a decrease in H3K27me3. MA plot showed total of 1063 regions that that are differentially enriched for changes in H3K27me3 in response to NOP16 overexpression (FDR<0.05). Compared to E.V. control, 755 regions showed a decrease while 308 regions showed an increase in H3K27me3.

**Extended Data Figure 4. NOP16 manipulation causes a misregulation of genes involved in adhesion, development, and tumorigenesis**

**a**. NOP16 overexpression causes a misregulation of genes involved in cell development and neurogenesis in MDA-MB231 cells. Dot plots represented gene ontology GO terms for all down-regulated genes in response to NOP16 stable overexpression as assessed by RNA sequence analysis. **b-c**. NOP16 deletion causes a misregulation of genes involved in morphogenesis, vascularization, and organization in MDA-MB231 cells. Dot plots represent categories of genes that are most significantly altered in gene expression as detected by RNA sequencing. **d**. Gene ontology enrichment map of down-regulated genes in MDA-MB231 cells in response to NOP16 knockout. **e.** Venn diagram of the overlap between NOP16 KO downregulated gene list with NOP16 overexpression upregulated gene list. Number of genes overlapping (20) occurs significantly higher than by random chance (p-value $< 1.0 \times 10^5$ as assessed by hypergeometric probability). Overlapping genes are presented on the right. **f.** Venn diagram reveals that no genes overlap between NOP16 KO upregulated and NOP16 OE downregulated gene lists. **g**. An unbiased gene set enrichment analysis of genes whose expression changes in response to NOP16 deletion showed highest similarity to genes that others have demonstrated to have high CpG levels in the promoter and are marked by H3K27me3.

**Extended Data Figure 5. Alteration of NOP16 expression affects the expression of gene sets involved in several tumor types as assessed by Disease GO term analysis**

**a.** Disease Ontology Enrichment of up-regulated genes in NOP16-stable overexpression compared with E.V. control. **b.** Heatmap of kidney cancer disease ontology enriched genes using lists of up-regulated genes in NOP16-stable overexpression compared with E.V. control. **c.** Higher expression of NOP16 correlates with poorer prognosis in kidney renal clear cell carcinoma patients. Plot is provided by Kaplan-Meier plotter. **d.** Disease Ontology Enrichment of up-regulated genes in NOP16 sgRNA compared n.g. control MDA-MB231 cells. **e.** Disease Ontology Enrichment of down-regulated genes in NOP16 sgRNA compared n.g. control MDA-MB231 cells. **f.** Heatmap of central nervous system cancer disease ontology enriched genes.

**Extended Data Figure 6. Several genes display changes in H3K27me3 that correlate with gene expression in response to NOP16 overexpression.**

Venn diagram of the overlap between NOP16 O/E upregulated gene list with decrease in H3K27me3 gene list using MDA-MB231 cells. This overlap (21 genes) is significantly higher than expected (p-value < 0.0001 as assessed by hypergeometric probability), and lower bar graph shows the GO analysis of these 21 overlap genes.

**Extended Data Figure 7. NOP16 knockdown decreases tumor cell growth *ex vivo* and *in vivo*.** **a.** Elevated expression of NOP16 correlates with an increasing frequency of genomic mutations, structural variants or copy number alterations to tumor-related genes. TCGA-BRCA datasets were obtained from cBioportal. **b.** NOP16 knockdown by siRNA is effective in breast cancer cells and mouse macrophage Raw264.7 cells as assessed by RT-qPCR. **c.** NOP16 knockdown in MCF7 cells decreases proliferation rates relative to a negative control (N/C) knockdown. $2.5 \times 10^4$ cells were plated and counted every other day. ****p<0.001 as assessed by two-way ANOVA followed by Dunnett's post-test. **d.** NOP16 knockdown in MDA-MB468 cells decreases proliferation rates relative to a negative control (N/C) knockdown. $5 \times 10^4$ cells were plated and counted every other day. ****p<0.001 as assessed by two-way ANOVA followed by Dunnett's post-test. **e.** *Nop16* knockdown in Raw264.7 cells suppressed cell growth relative to a negative control (N/C) knockdown. $5 \times 10^4$ cells were plated and counted every day. ****p<0.001 as assessed by two-way ANOVA followed by Dunnett's post-test. **f and g.** CRISPR-mediated knockout of NOP16 is efficient in MCF7 (f) and MDA-MB231 (g) cells as assessed by western blotting. **h.** NOP16 knockout slows cell proliferation in MDA-MB231 cells. MDA-MB231 cells were infected with lentivirus encoding Cas9 and single guide RNAs directed against control or NOP16 (#1 or #4), after selection and recovery $2.5 \times 10^4$ cells were seeded and cell numbers were counted every other day. ****p<0.001 as assessed by two-way ANOVA followed by Dunnett's post-test. **i.** A heat map reveals decrease in cell cycle genes in response to NOP16 depletion in MDA-MB231 cells. **j.** NOP16 is efficiently knocked-down by siRNA in HeLa-Fucci(CA5) cells as assessed by RT-qPCR. **k and l.** NOP16-AsiC treatment decreased the levels of NOP16 mRNA and the cell viability in MDA-MB468 cells *ex vivo*. MDA-MB468 cells were treated with NOP16 AsiC (1, 2, 4 μM) and Aptamer control (4 μM) for 72 hrs. i) The total RNA extracted from each sample and NOP16 mRNA expression were determine by RT-qPCR normalized to GAPDH. j) The cell viability was evaluated by Cell-titer Glo. *p<0.05, **p<0.01, ***p<0.001, ****p<0.0001 as assessed by one-way ANOVA followed by Dunnett's post-test. **m.** NOP16-AsiC treatment caused d a decrease in NOP16 expression in MDA-MB468 induced tumor tissues from mice. NOP16 expression was evaluated by RT-qPCR and normalized to GAPDH. **p<0.01 as assessed by student t-tests.

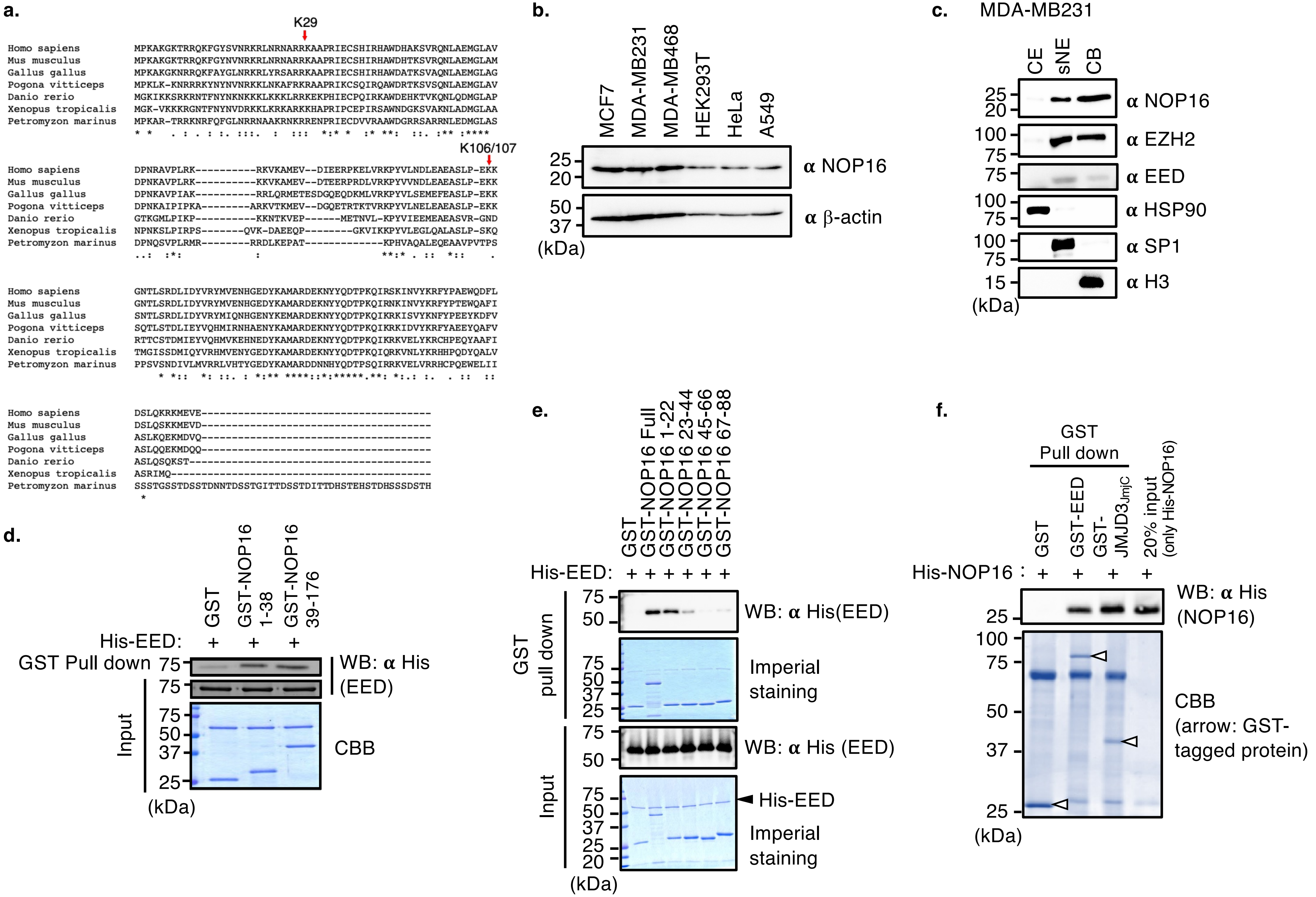


Takashima K et al Extended Data Figure 1

**a.**

KPYVLNDLEAEASLPEKKGNTL

**b.**

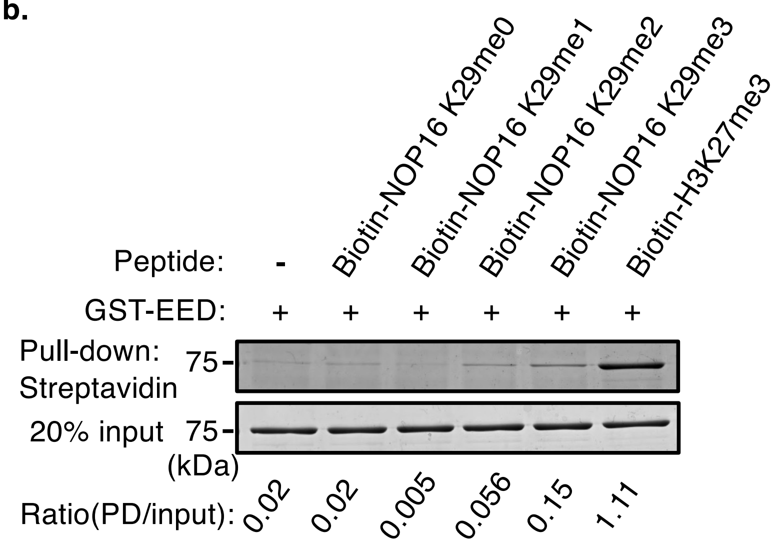


**c.**

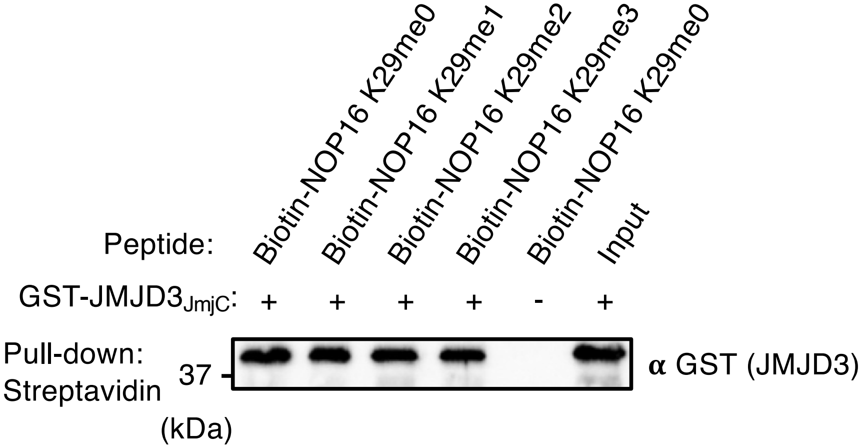


**d.**

| Peptide | $K_D$(μM) ± STDV |
|---|---|
| H3K27me3 | 8.59 ± 4.54 |
| NOP16K29me3 | 32.15 ± 27.37 |
| NOP16K29me2 | 305.6 ± 100.8 |
| NOP16K29me1 | 346.67 ± 224.26 |
| NOP16K29me0 | 102.68 ± 65.25 |

**a.** MDA-MB231

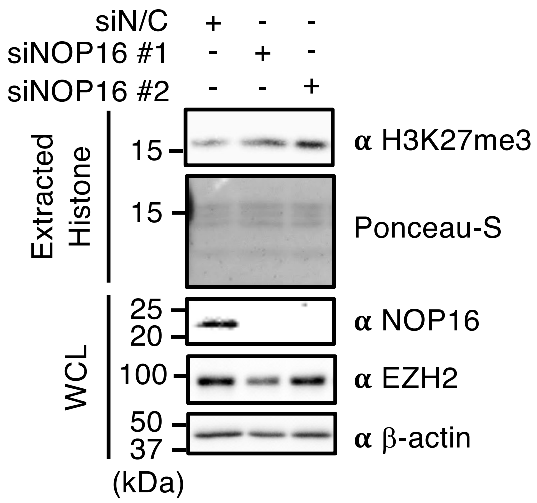


**b.** CUT&RUN

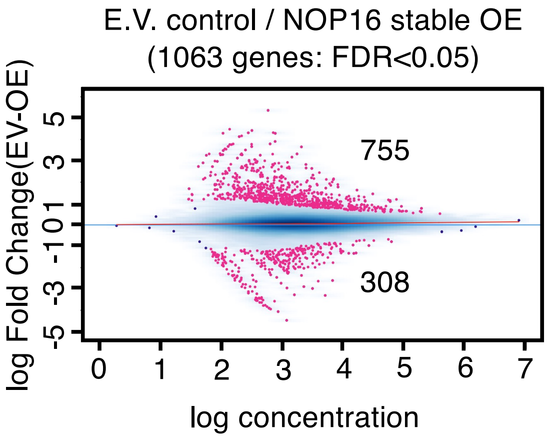

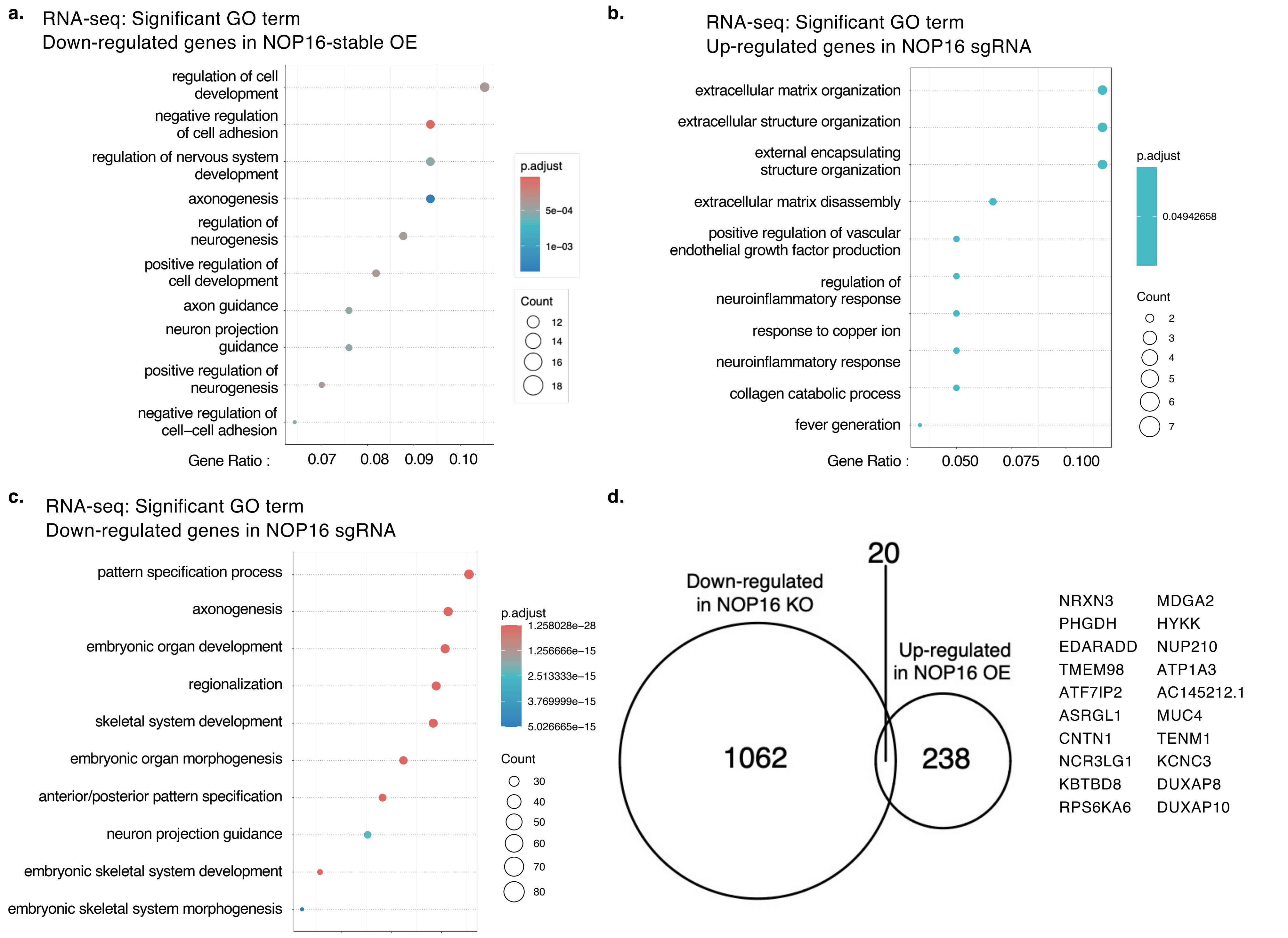


Takashima K et al Extended Data Figure 4

e.

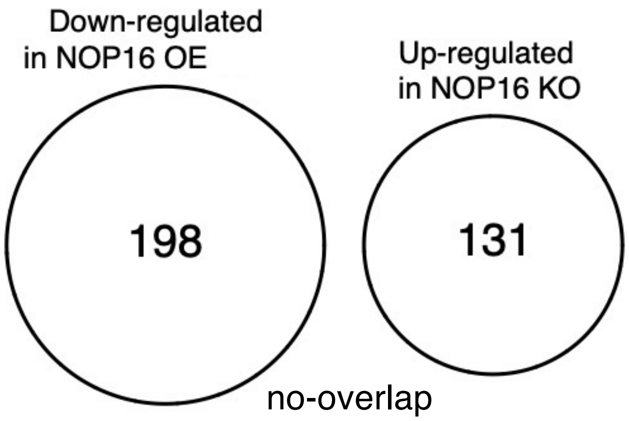


f.

| RNA-seq : MSigDB Curated gene sets(C2): ng vs NOP16 KO | NES | p.adjust |
|---|---|---|
| MEISSNER_BRAIN_HCP_WITH_H3K27ME3 | -1.784158 | <1.00E-6 |
| MEISSNER_NPC_HCP_WITH_H3K27ME3 | -1.861147 | <1.00E-6 |
| MEISSNER_NPC_HCP_WITH_H3K4ME2 | -1.58223 | <1.00E-6 |
| MEISSNER_NPC_HCP_WITH_H3K4ME2_AND_H3K27ME3 | -1.712808 | <1.00E-6 |
| MIKKELSEN_IPS_WITH_HCP_H3K27ME3 | -1.837471 | <1.00E-6 |
| MIKKELSEN_MCV6_HCP_WITH_H3K27ME3 | -1.735195 | <1.00E-6 |
| MIKKELSEN_MEF_HCP_WITH_H3K27ME3 | -1.697657 | <1.00E-6 |
| MIKKELSEN_NPC_HCP_WITH_H3K27ME3 | -1.775413 | <1.00E-6 |
| REACTOME_MITOCHONDRIAL_TRANSLATION | 2.405705 | <1.00E-6 |
| REACTOME_NEURONAL_SYSTEM | -1.581208 | <1.00E-6 |
| MIKKELSEN_NPC_HCP_WITH_H3K4ME3_AND_H3K27ME3 | -1.65832 | <1.00E-6 |
| SABATES_COLORECTAL_ADENOMA_DN | -1.580479 | 0.000001 |
| MEISSNER_NPC_HCP_WITH_H3K4ME3_AND_H3K27ME3 | -1.704845 | 0.000001 |
| DAWSON_METHYLATED_IN_LYMPHOMA_TCL1 | -1.8461 | 0.000001 |
| LIU_PROSTATE_CANCER_DN | -1.461872 | 0.000001 |
| LIM_MAMMARY_STEM_CELL_UP | -1.461482 | 0.000003 |
| NAKAYAMA_SOFT_TISSUE_TUMORS_PCA1_DN | -1.796087 | 0.000004 |
| REACTOME_EUKARYOTIC_TRANSLATION_INITIATION | 2.171445 | 0.000004 |
| VERHAAK_GLIOBLASTOMA_PRONEURAL | -1.624144 | 0.000005 |

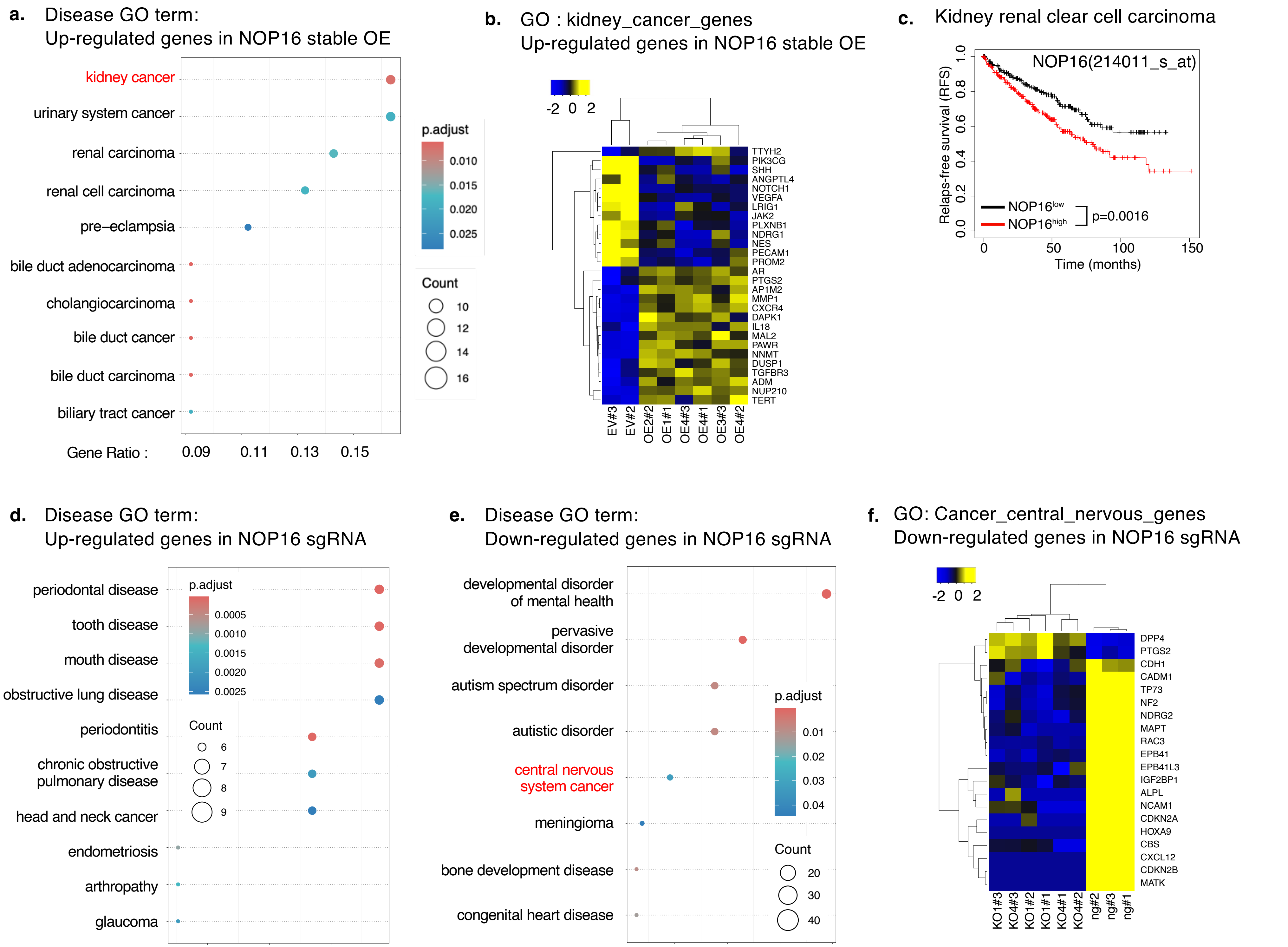


Takashima K et al Extended Data Figure 5

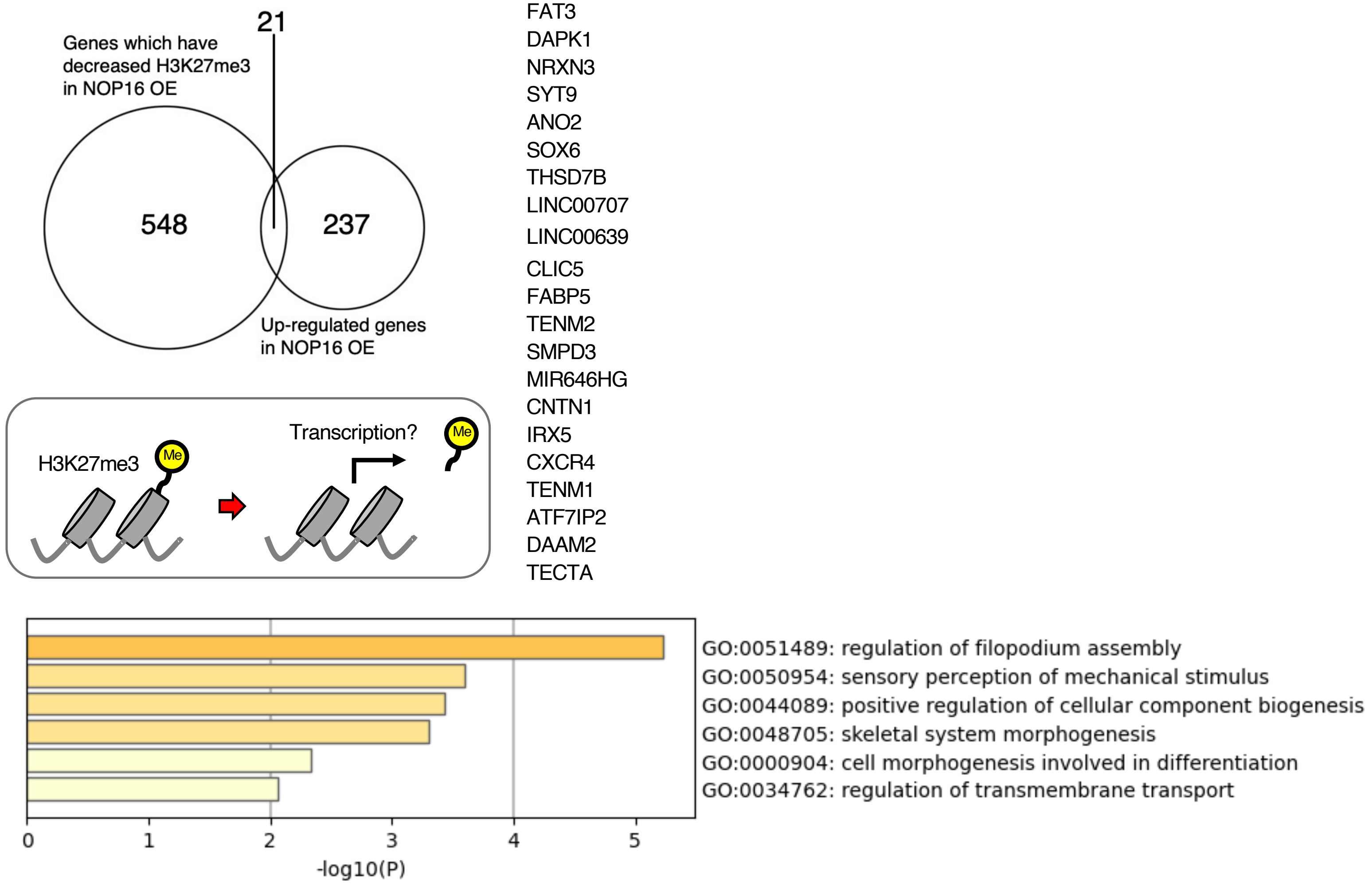


Takashima K et al Extended Data Figure 6

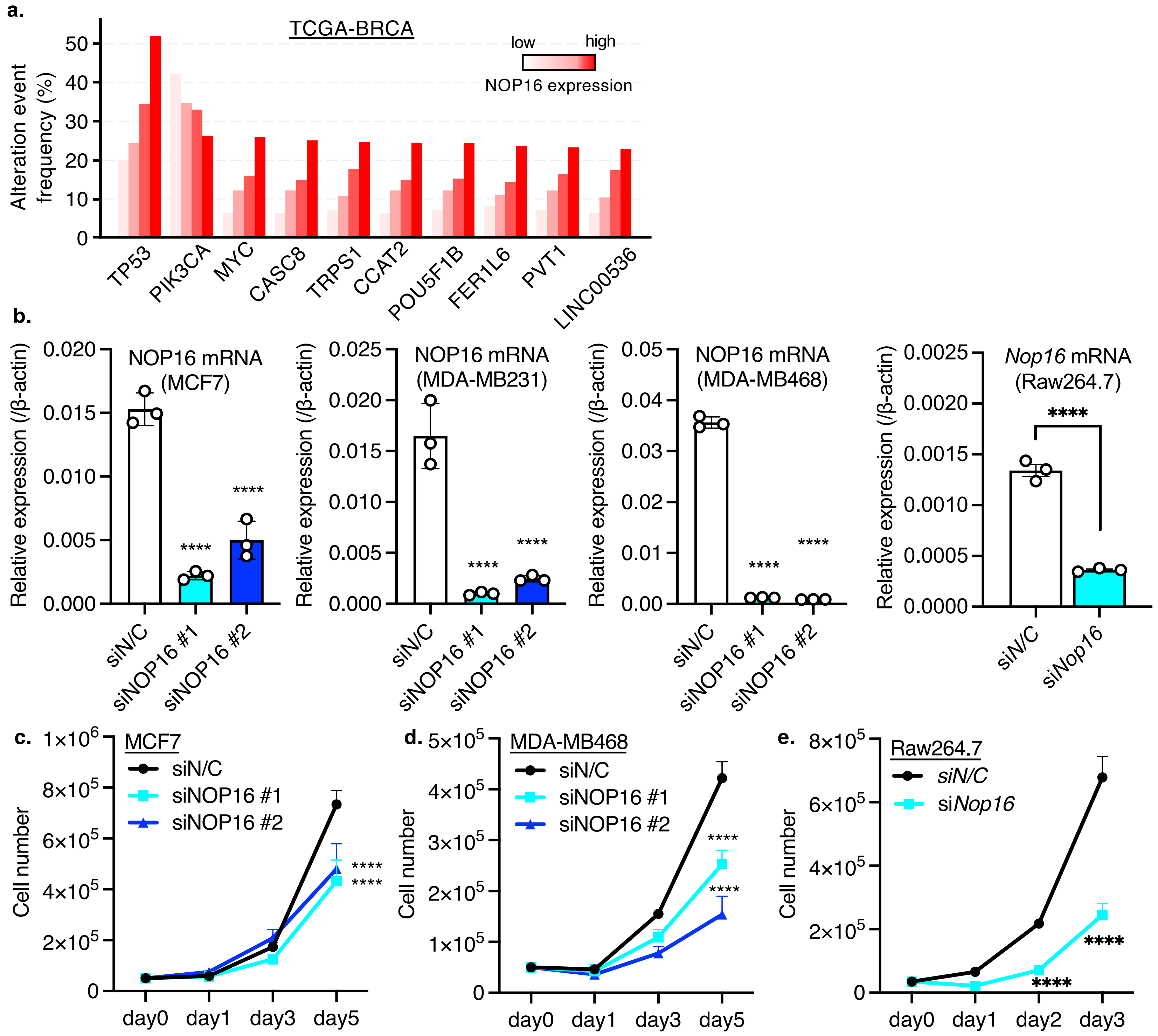


Takashima K et al Extended Data Figure 7

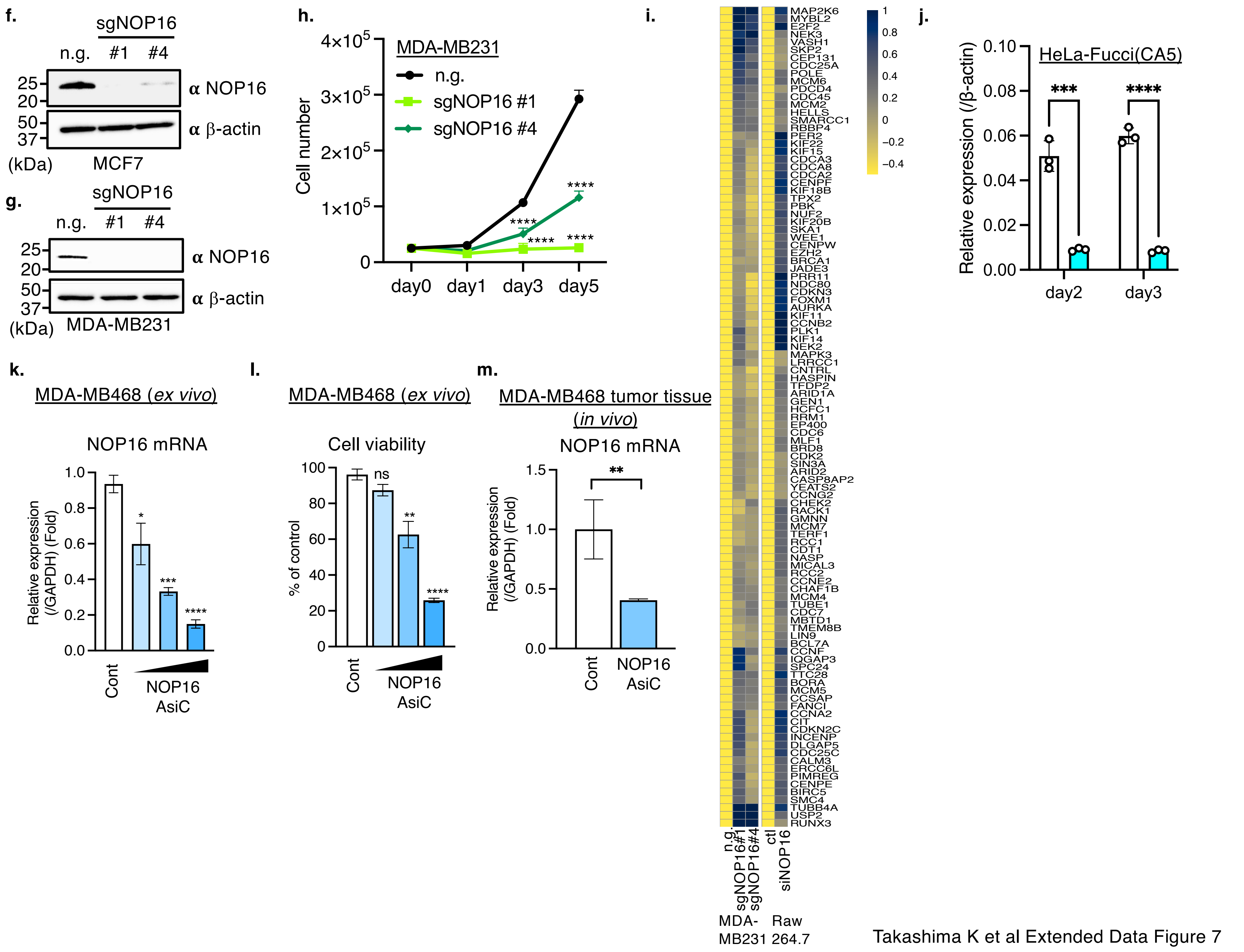


Takashima K et al Extended Data Figure 7
